# Functional imaging of conduction dynamics in cortical and spinal axons

**DOI:** 10.1101/2023.02.28.530461

**Authors:** Milos Radivojevic, Anna Rostedt Punga

## Abstract

Mammalian axons are specialized for transmitting action potentials to targets within the central and peripheral nervous system. A growing body of evidence suggests that, besides signal conduction, axons play essential roles in neural information processing, and their malfunctions are common hallmarks of neurodegenerative diseases. The technologies available to study axonal function and structure integrally limit the comprehension of axon neurobiology. High-density microelectrode arrays (HD-MEAs) allow for accessing axonal action potentials at high spatiotemporal resolution, but provide no insights on axonal morphology. Here we demonstrate a method for electrical visualization of axonal morphologies based on extracellular action potentials recorded from cortical and motor neurons using HD-MEAs. The method enabled us to reconstruct up to 5-centimeter-long axonal arbors and directly monitor axonal conduction across thousands of recording sites. We reconstructed 1.86 meters of cortical and spinal axons in total and found specific features in their structure and function.

## INTRODUCTION

Axons are neuronal processes specialized for the conduction of action potentials (APs). Cortical axons serve as communication cables between various types of neurons that are synaptically connected and, accordingly, arranged in multiple layers to receive, process, and convey neural information between different regions in the brain. Motor neurons located in the ventral horn of the spinal cord project their axons outside the central nervous system (CNS) and are responsible for the contraction of effector muscles in the periphery. Spinal axons are specialized to innervate and precisely control different types of muscle fibers, thus ensuring refined coordination of complex body-movements^1^.

Mainly due to difficulties to experimentally access axonal conduction, axonal information processing has been neglected, and axons are classically seen as conductive cables that do nothing more than faithfully transmit APs in an all-or-none, ‘binary’ fashion^2^. Later studies have challenged this view and suggested that axons have much more complex roles than traditionally thought^3^. Contrary to classical concepts, reports show that, besides conducting binary APs, hippocampal and cortical axons can also transmit analog currents in a passive fashion^4, 5^. It has been shown that analog currents integrate with ongoing axonal APs, changing their waveforms and, consequently, affecting synaptic transmission in a graded, ‘analog’ manner^4, 6, 7^. Additionally, it was found that axons modulate the waveforms of APs contingent on neuronal activity and, accordingly, adjust the synaptic release^8^. Moreover, during high-frequency regimes of neuronal activity, axons are able to reduce their conduction velocities by more than 20% and, thereby, tune the timing of AP arrival at synapses^9^. Taken together, these findings suggest that axons passively and actively process APs to tune the amount of information transmitted by synapses, but also imply that axons play a crucial role in the temporal coding of the neural information.

Morphological complexity of mammalian axons generally depends on their downstream targets’ spatial disposition. Thus, for example, cortical neurons form extensively branched axons to convey APs to numerous postsynaptic neurons located in different cortical layers and regions in the brain. Due to their small diameter of 0.08-0.4 µm^10^ and absence of surrounding myelin sheaths, cortical axons provide relatively low conduction velocities of 0.1-2 m/s^9, 11^, which are, nevertheless, sufficient to ensure rapid communication between closely spaced neurons. As opposed to cortical neurons, motor neurons transmit signals to distant targets outside the CNS and, for that purpose, develop considerably longer (up to 1 m), thicker (0.5-10 µm), and myelinated axons that provide considerably higher conduction velocities of up to 100 m/s^12, 13^.

Despite being acknowledged as reliable conduction cables, axons can in certain cases fail to propagate APs faithfully. Such cases are referred to as conduction failures and are attributed to particular axonal morphology. For instance, experimental and theoretical studies suggest conduction failures are likely to occur at axonal branching points and local swellings due to abruptly increased axon diameter^10^. In addition, the AP propagation fidelity and temporal precision depend on axon diameter, biophysical properties of various types of ion channels and thermodynamic noise inherent to their gating dynamics^14–16^. According to theoretical studies, thin distal axons (diameter < 1 µm) are prone to “channel noise”, which can introduce variability in axonal AP waveforms^17^, increase the jitter of AP propagation^9, 14^, and compromise the reliability of the conduction itself^14, 18, 19^. Besides geometrical factors, neuronal activity can also affect AP conduction, which is consistent with our previous finding that high-frequency neuronal activity increases the jitter of AP propagation in cortical axons^9^.

Ultrastructural aberrations and malfunctioning conduction in human axons are often early hallmarks of neurodegenerative diseases. For instance, in Alzheimer’s disease (AD), abnormal protein aggregates cause local swellings in axons which have been shown to reduce conduction velocity and even block AP propagation^20^. Human post-mortem studies suggest axonal degeneration may be the earliest feature of Parkinson’s disease (PD) and, therefore, an appropriate target for early therapeutic interventions^21^. Motor neuron degeneration is the hallmark of amyotrophic lateral sclerosis (ALS), where axonal dysfunction begins long before symptom onset and motor neuron death^22^.

Comprehension of axon neurobiology in health and disease is generally limited by the technologies available to study axonal function and structure integrally. The whole-cell patch-clamp technique, complemented with fluorescence microscopy, is commonly used to correlate electrophysiological data with the morphological properties of the neuron. Namely, a patch-clamp pipette can inject fluorescent dyes directly into the neuron and, therefore, allow visualization of the axonal arbor during the electrophysiological experiment. However, patch-clamp is typically limited to recording APs from a single axonal site^4, 6, 23^ and does not allow for tracking AP propagation across axons. Moreover, the technique is invasive and destructive, which constrains the duration of a recording session to about an hour. Alternatively, fluorescent indicators sensitive to voltage^24^ or calcium^25^ can be used to observe axonal AP propagation and, at the same time, to visualize axonal morphology. However, fluorescent indicators exhibit photobleaching and phototoxicity and may perturb the physiology of the cell to the point of affecting AP conduction^24^. Complementary-metal-oxide-semiconductor (CMOS) -based high-density microelectrode arrays (HD-MEAs) have been designed to record extracellular APs from neuronal cultures^26^ and allow tracking axonal signals across hundreds of microelectrodes^9, 11^. Thanks to a low-noise CMOS-design, HD-MEAs enable detection of APs across entire arbors of cortical axons, including tiny axon terminals^9^. HD-MEAs provide noninvasive access to axonal APs and impose no constraints on the duration of the recording sessions. Nevertheless, HD-MEA technology does not provide direct insights into axonal morphology. It has to be complemented with auxiliary optical techniques to allow correlation between axonal function and structure. Live-cell imaging techniques can be used to optically visualize axonal morphologies directly on HD-MEA surfaces^9, 11, 27^. These techniques, however, entail introduction of fluorescent reporters into the cell, which induces chemotoxicity, phototoxicity and cell death^28^.

Our objective was to enable the reconstruction of axonal morphologies based solely on extracellular APs recorded from *in vitro* mammalian neurons using the CMOS-based HD-MEA system. The present study reports a method for tracking AP propagation across tens-of-millimeter-long nonmyelinated axonal arbors of primary cortical and motor neurons cultured on HD-MEAs. The method allows for label-free electrical visualization of axonal conduction trajectories and, at the same time, provides noninvasive access to axonal APs waveforms recorded across hundreds of microelectrodes. Using the developed method, we investigated (I) morphological features of cortical and spinal axons, (II) fluctuations in AP amplitudes across axonal arbors, and (III) temporal dynamics of the axonal conduction.

## RESULTS

We developed a method for the automatic reconstruction of axonal morphology based on extracellular APs recorded from cortical and motor neurons using the HD-MEA system (Fig. 1A). The reconstructed morphology is here referred to as “functional morphology”, and it reveals the location of the axon initial segment (AIS), axonal trunk and higher order branches (Fig. 1B). The method was used to trace extracellular APs propagating across axonal arbors of cortical (Movie S1) and motor neurons (Movie S2). The experimental design and overview of the method are outlined in Fig. 2 and are described below.

**Figure 1.**
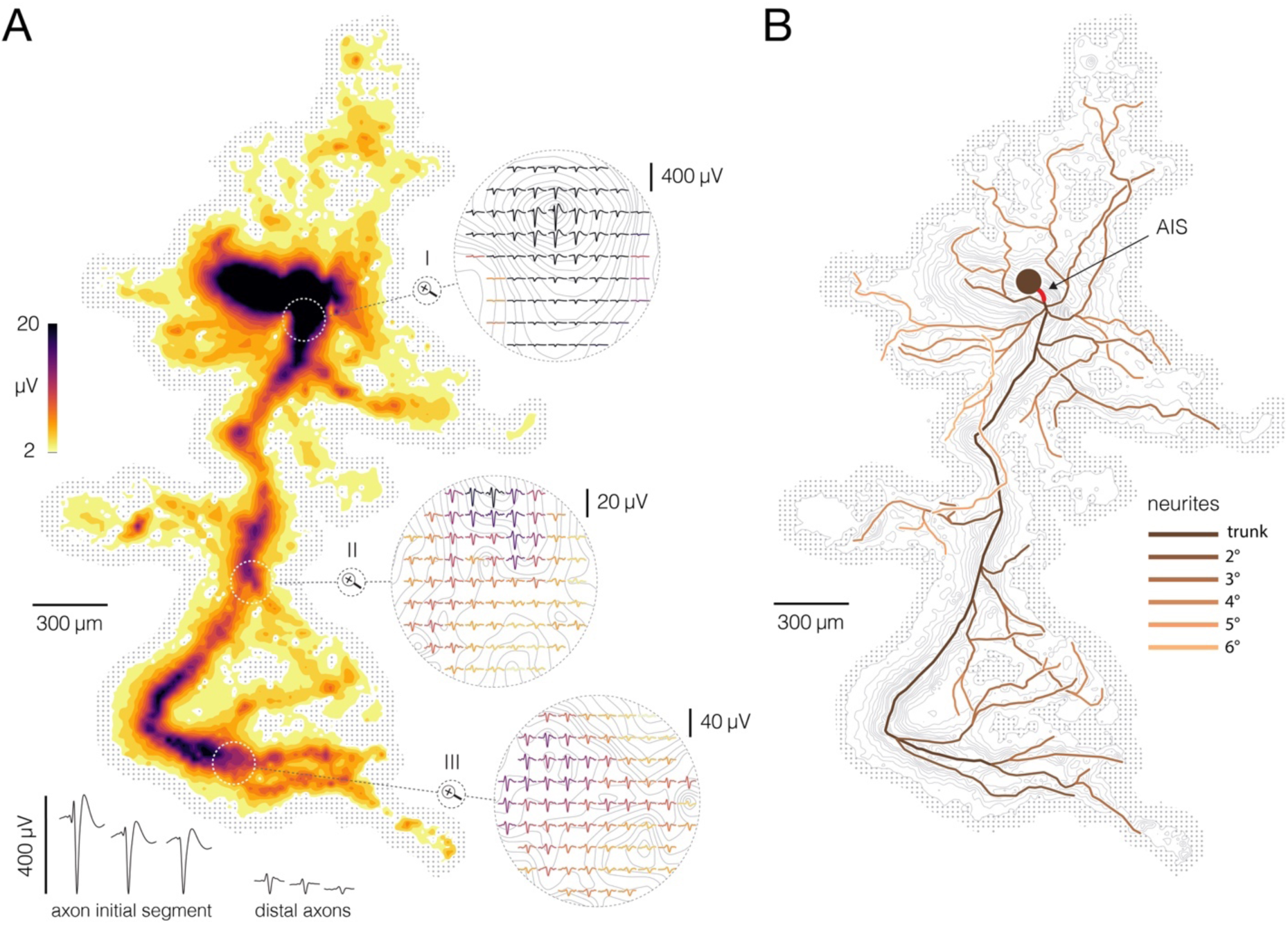
Reconstruction of axonal morphology based on neuronal electrical activity. (**A**) Contour map shows spatial distribution of extracellular action potentials (APs) recorded from a single cortical neuron. Average AP amplitudes are color-coded and presented through contour surfaces. Grey dots in the background represent locations of the recording electrodes. White-dashed-line circles superimposed over the contour map denote three magnified regions displayed on the right. Average AP waveforms obtained from proximal, middle and distal axons are shown in the three denoted regions (labelled I, II and III, respectively); color-coding is the same as in the contour map. Examples of AP waveforms recorded from the axon initial segment (AIS) and distal axons are shown at the bottom. (**B**) Functional morphology of axonal arbor reconstructed from APs displayed in (A). Branching orders of reconstructed neurites are color-coded. The AIS is depicted by a thick red line superimposed over the most proximal part of the axon. A filled dark-brown circle represents putative location of the neuronal soma. Color-free contour map and recording electrodes are shown in the background (same as in A). Functional morphologies of cortical and spinal axons are also presented in Movie S1 and Movie S2, respectively.

**Figure 2.**
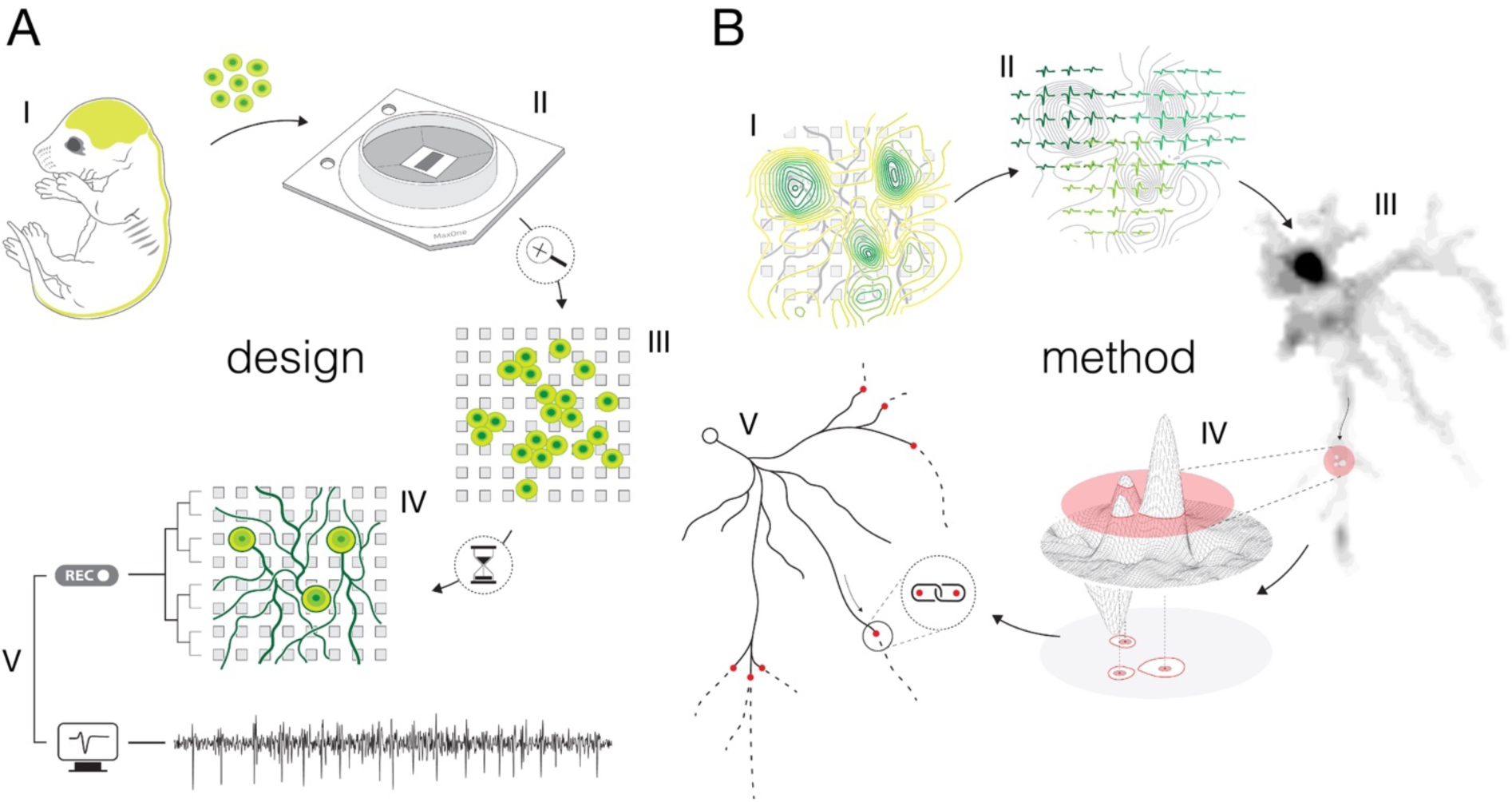
Experimental design and overview of the method. **(A)** Experimental design: (I) Brain cortices and spinal cords isolated (lime-green) from Sprague Dawley rat embryos were used as sources of primary cortical and motor neurons. Cells obtained from corresponding sources were seeded directly on HD-MEAs to grow cortical and motor neurons in separate cultures. (II) The utilized HD-MEA system features 26,400 microelectrodes packed within an area of 3.85×2.1 mm^2^. Interelectrode distances of 17.5-µm provide density of 3,264 microelectrodes per mm^2^. (III) The HD-MEA surface coated with cell-adhesive materials promoted neuronal growth over the sensing area and provided efficient electrode-to-neuron contacts. (IV) Dedicated media and growth-promoting factors were used to develop and maintain the cultures *in vitro* over extended periods of time. (V). Mature cultures conveyed spontaneous electrical activities. Neuronal extracellular APs were recorded through HD-MEA recording channels at 20 kHz sampling rate. **(B)** Overview of the method: (I) Signal amplitudes sampled across an entire HD-MEA were used to produce network-wide activity maps (yellow-green contour map). Local peaks found in activity maps indicated locations of individual neurons in the network. (II) Spike-sorting of recorded signals enabled to discern activities between neighboring neurons and to extract their individual electrical profiles. Displayed electrical profiles (dark-, mint- and lime-green waveforms) are referred to as “extracellular AP footprints”. (III) Array-wide spike-triggered averaging revealed spatiotemporal distribution of APs across axons of individual neurons (grey-scale signal map). Such representation of neuronal signals is referred to as an “axonal electrical image”. (IV) Dynamic thresholding was used to detect local peaks within the footprint. (V) Detected peaks were interlinked based on their spatial and temporal proximities to reconstruct axonal conduction trajectories.

### Experimental design and overview of the method

We cultured rat primary cortical (Fig. S1) and motor neurons (Fig. S2) on a CMOS-based HD-MEA system that comprises 26,400 densely packed microelectrodes (Fig. S3). The cultures were grown in monolayers with estimated thicknesses of ∼5-40 and ∼5-50 µm for cortical and motor neuron cultures, respectively (Fig. S4). Both cortical and motor neuron cultures yielded cell-densities of ∼500-2,000 neurons per mm^2^ (Fig. S4). The cultures matured in controlled conditions (see methods), and their extracellular electrical activity was recorded between 12 and 24 days in vitro (DIV). Mature neurons grew their axonal arbors efficiently across sensing areas of the HD-MEA chips and provided a tight interface between axons and microelectrodes (Fig. S5). The relative proximity of axons to the sensing area varied across axonal arbors within ranges of ∼1-13 and ∼1-18 µm for cortical and motor neurons, respectively (Fig. S5).

The CMOS-based HD-MEA system enabled us to map extracellular APs across entire cultures and to electrically identify individual neurons (Fig. S6). Spontaneous neuronal activities were recorded across all microelectrodes for 2 minutes, and average amplitudes of the recorded voltage traces were used to produce network-wide activity maps (Fig. S6A). Because extracellular APs with the largest amplitudes arise from the AIS^29^, local maxima found within these maps indicated the location of individual neurons in the network^30^.

We used spike-sorting algorithms to discern signals among individual neurons in the culture (see methods). The dense arrangement of the microelectrodes allowed us to access the electrical activity of a single neuron at high spatial resolution (Fig. S6A, Movie S3, S4); however, overlapping signals recorded from multiple neighboring neurons were observed in most of the cases (Fig. S6B). Spike-sorting procedures enabled us to discern APs among individual neurons reliably and extract relative times of their activities (“spike times”)^9, 30, 31^. Sorted APs were then averaged over adjacent electrodes to reveal the spatiotemporal distribution of a single neuron’s activity. The spatiotemporal distribution of APs recorded from proximal neuronal compartments (near the AIS) is called an “extracellular AP footprint”. Electrical footprints reconstructed for eight neighboring neurons are presented in Fig. S6C. Z-stack image series of the corresponding culture is shown in Movie S3, S4.

We used spike-triggered averaging for electrical imaging of axonal arbors (Fig. S7). High-amplitude APs detected near the AIS were used to trigger the averaging of single voltage traces (‘single trials’) recorded across all electrodes in the array (Fig. S7A). Because the AIS signal represents the first occurring (initial) trace of the neuron’s activity, the averaging reveals spatial and temporal shifts in propagating axonal signals (Fig. S7B). The spatiotemporal distribution of axonal signals reconstructed using spike-trigger averaging is called an “axonal electrical image”. Axonal electrical images of three neighboring neurons are presented in Fig. S7B and in Movie S5.

We developed a method for reconstructing axonal functional morphologies based purely on features extracted from axonal electrical images. Adaptive thresholding was used to map signal peaks across axonal arbors (Fig. 3), and the moving object tracking technique was used to reconstruct axonal conduction trajectories (Fig. 4). The method for reconstruction of axonal functional morphologies is described in the following two sections.

**Figure 3.**
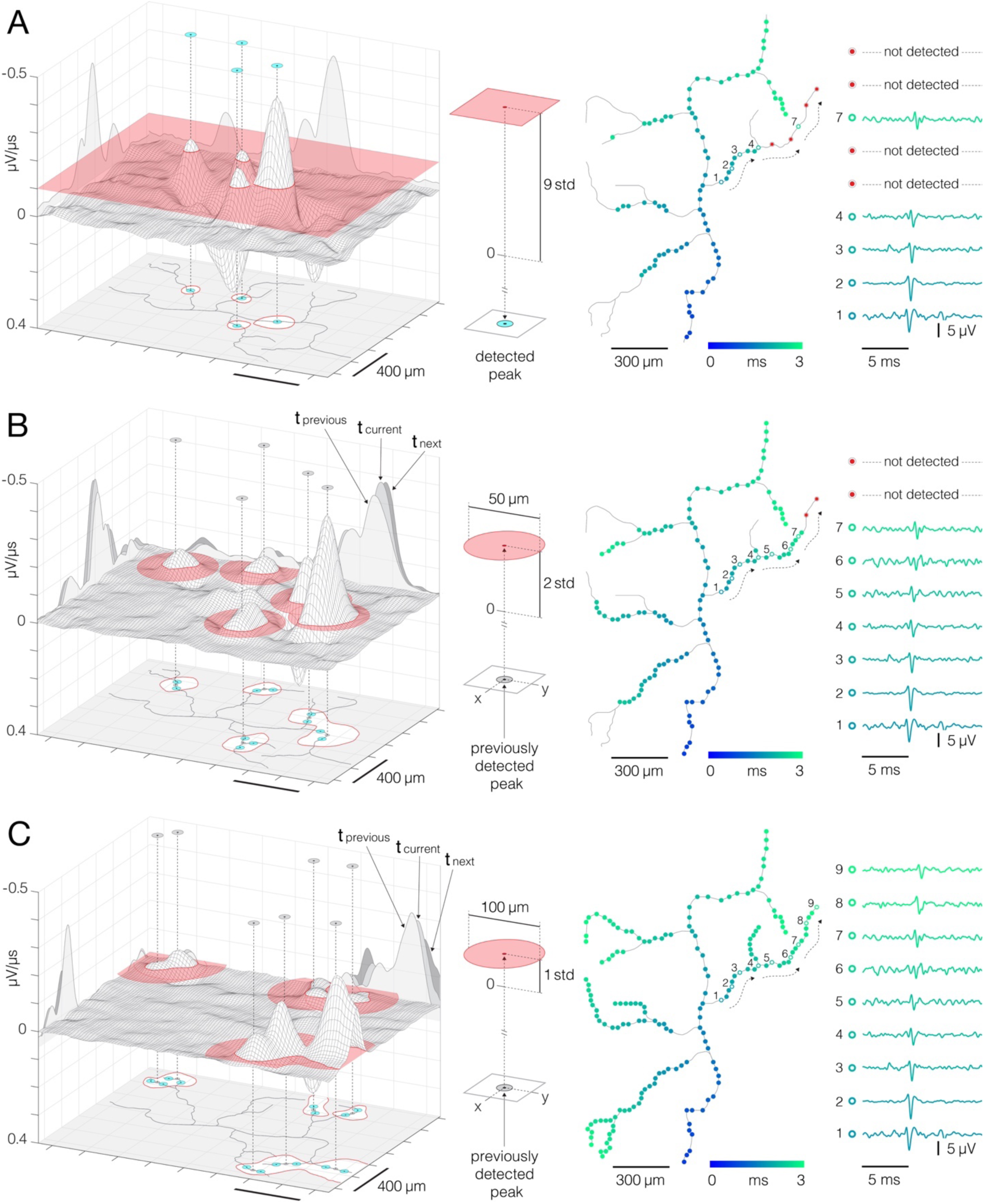
Detection of action potentials propagating across axonal arbors. **(A)** A simple threshold enables detection of high-amplitude signal peaks. (Left) 3D graph shows planar threshold (red semitransparent plane) applied to a section of neuronal signal (3D gray-lined mesh) reconstructed for a single timeframe. 2D profiles of the reconstructed signal are projected on the graph’s side planes (gray-filled hills). Signal cutouts found above the threshold are projected on the graph’s bottom (red-line bordered white patches). Detected signal peaks are denoted by cyan circles projected perpendicularly from top to the bottom of the graph. Axonal contour shown at the graph’s bottom was estimated by observing spatial movements of the signal peaks over consecutive timeframes (also see Movie S12, S13). (Middle) Simple planar threshold was set to 9 STD of the estimated noise; background noise was sampled across all electrodes during periods when the neuron was inactive. (Right) Spatiotemporal distribution of the detected AP peaks (blue-cyan circles) superimposed over axonal contour (same as in the left panel). Timing of the detected peaks is color-coded. Action potential waveforms recorded at the numbered locations in the upper-right axonal branch are shown on the side. Black-dashed arrows indicate direction of the propagating APs. Red circles denote signal peaks that were not detected by the simple threshold. **(B, C)** Adaptations of the threshold based on spatial and temporal coordinates of previously detected peaks enabled detection of low-amplitude signals. (Left) Locally applied thresholds (red semitransparent circles) were centered on XY coordinates of previously detected peaks (gray circles projected perpendicularly from top to the bottom of the graph); local thresholds are adapted to detect neighboring peaks at preceding and succeeding timeframes. Neuronal electrical activity reconstructed for a single timeframe (3D gray-lined mesh) is projected on the graph’s side planes (t_current_); signal profiles of preceding and succeeding timeframes are also projected (t_previous_, t_next_). Signal cutouts found above the threshold are projected on the graph’s bottom (red-line bordered white patches). Newly detected peaks are denoted by cyan circles projected on the graph’s bottom plane. Axonal contour same as in (a). (Middle) Detection fields of local thresholds are 50 µm (in B) and 100 µm (in C) in radius, and are set to 2 STD (in B) and 1 STD (in C) of the estimated noise. (Right) Same as in (A); note newly mapped peaks detected near proximal axon. Peak detection strategies are comprehensively demonstrated in Movie S6-8.

**Figure 4.**
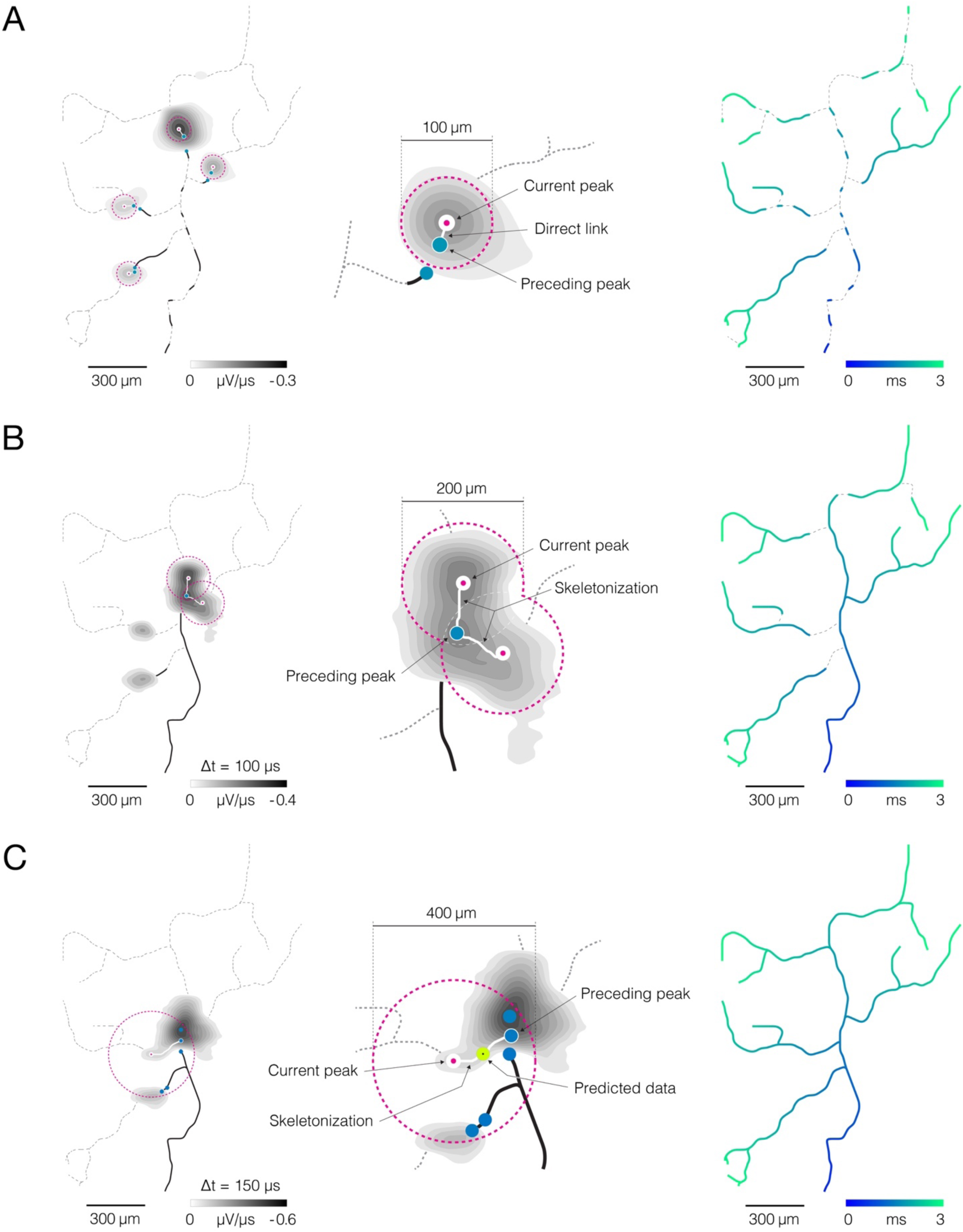
Reconstruction of axonal functional morphology. **(A)** Direct interconnection of mapped AP peaks. (left-middle) Closest signal peaks found within a 100-µm Euclidean range (pink-dashed circles) and mapped in two consecutive timeframes (current and preceding peak) were interconnected via direct links (rectilinear white lines). Axonal APs reconstructed for the current timeframe is shown in the background (contour map); average signal amplitude is color-coded (gray scale). (right) Axonal conduction trajectories revealed by direct interconnection; timing of the propagating AP is color-coded (blue-cyan). **(B)** Skeletonization-assisted interconnection of mapped signal peaks. (left-middle) Signal peaks found within a 200-µm Euclidean range (pink-dashed circles) and mapped at two consecutive timeframes (current and preceding peak) are interconnected via remnants of the skeletonized signal (irregular white lines) - signal averaged over the two consecutive timeframes (Δt = 100 µs) is skeletonized to infer directionality of the propagating signal. Axonal signal averaged over the two consecutive timeframes is shown in the background (contour map); signal amplitude is color-coded (gray scale). (right) Axonal conduction trajectories reconstructed using direct and skeletonization-assisted interconnection; timing of the propagating signal is color-coded (blue-cyan). **(C)** Indirect interconnection of mapped signal peaks. (left-middle) Signal peaks that were mapped at every other timeframe and found within a 400 µm Euclidean range (pink-dashed circles) were interconnected indirectly – the signal averaged over three consecutive timeframes (Δt = 150 µs) was skeletonized to interconnect peaks mapped in 1^st^ and 3^rd^ timeframe (irregular white line); data for the 2^nd^ timeframe was predicted based on the conduction velocity observed in previously reconstructed trajectories. Axonal signal averaged over the three consecutive timeframes is shown in the background (contour map); signal amplitude is color-coded (gray scale). (right) Functional axonal morphology reconstructed using direct, skeletonization-assisted and indirect interconnection; timing of the propagating APs is color-coded (blue-cyan). Direct, skeletonization-assisted and indirect interconnections of mapped signal peaks are demonstrated in Movie S9-11.

### Adaptive thresholding enables mapping of axonal signal peaks in space and time

Adaptive thresholding applied to axonal electrical images enabled us to map AP peaks propagating across axonal arbors (Fig. 3). Axonal electrical images were obtained by averaging 200 voltage traces per electrode. Time derivatives of averaged traces (µV/µs) were computed for each of the electrodes, and the resulting data were divided into 400 consecutive timeframes (with 50 µs inter-frame intervals). Adaptive thresholds were next applied at each timeframe to detect AP peaks at different time points during axonal propagation.

Our thresholding technique generally utilized a ‘greedy algorithm’ principle (see discussion). The algorithm allowed for recurrent adaptations of the threshold based on parameters updated along the detection process. High-amplitude APs were mapped first, and parameters obtained from the mapped locations were used to tailor the detection of low-amplitude signals in the later steps. The thresholds were initially determined based on electrical noise observed from the HD-MEA chip. Electrical noise was estimated from voltage traces sampled across an entire array during periods when the observed neuron was inactive, and the noise was estimated for each neuron separately.

Adaptive thresholding involves three steps (Fig. 3, Movie S6-8). In the first step, a simple planar threshold, set to 9 STD of the estimated noise, was used to detect high-amplitude signal peaks (Fig. 3A, Movie S6). A high threshold level allowed us to detect AP peaks far above the background noise. However, axonal APs with lower amplitudes remained undetected, leaving gaps along axonal conduction trajectories. In the second step, confined thresholds, set to 2 STD of the estimated noise, were applied locally to detect low-amplitude AP peaks (Fig. 3B, Movie S7). The confined thresholds were positioned on spatial and temporal coordinates of previously mapped peaks. The thresholds were confined spatially to 50 µm radii and temporally to periods encompassing three consecutive timeframes (t_previous_, t_current_, t_next_, see Fig. 3B). The spatiotemporal confinement enabled us to detect low-amplitude signals in close proximity to previously mapped peaks and to fill local gaps along the conduction trajectories. The third step utilized the same detection strategy but with differently tuned parameters (Fig. 3C, Movie S8). Namely, the threshold level was further lowered to 1 STD of the estimated noise, and the spatial confinement was broadened to a 100 µm radius. These parameters enabled the detection of low-amplitude APs near axon terminals.

### A moving object tracking technique enables reconstruction of axonal conduction trajectories

We developed an algorithm for tracking mapped axonal signals across consecutive timeframes (Fig. 4). The algorithm enabled us to reveal signal conduction trajectories and reconstruct functional morphologies of cortical (Movie S1) and spinal axons (Movie S2).

Our tracking algorithm was designed to predict axonal conduction trajectories based on three factors: (I) spatiotemporal proximities of the mapped peaks, (II) topology of skeletonized axonal signal, and (III) conduction velocities estimated from previously reconstructed trajectories.

The tracking procedure involved three iterative steps (Fig. 4, Movie S9-11). (I) Direct interconnection – the closest signal peaks found within a 100 µm Euclidean distance and mapped in two consecutive timeframes were interconnected via direct links (Fig. 4A, Movie S9). Direct interconnection revealed fragments of axonal conduction trajectories but failed to reconstruct axonal branching forks in most cases. Reconstructed fragments were used to calculate axonal conduction velocities. Obtained velocities served as criteria to tailor the tracking procedure in the next two steps. (II) Skeletonization-assisted interconnection – consecutive signal peaks that could not be interconnected directly, but were found within a 200 µm Euclidean range, were interconnected via skeletonized remnants of the axonal signal (Fig. 4B, Movie S10). Signal amplitudes extracted from the corresponding electrical images were averaged over two consecutive timeframes (Δt = 100 µs) and mapped over interconnection areas. Obtained maps were next skeletonized to infer propagation trajectory between the consecutive signal peaks. Conduction velocities estimated in the previous step were used as criteria for the selection of optimal propagation trajectories. Trajectories whose velocities deviated from previously estimated values by more than 50 % were discarded. Skeletonization-assisted interconnection enabled reconstruction of some, but not all, axonal branching forks. It was designed to predict axonal trajectories over consecutive (continuous) signal peaks but could not predict trajectories between discontinuous signal peaks. Axonal conduction velocities estimated from the reconstructed trajectories served as criteria to tailor the tracking procedure in the next step. (III) Indirect interconnection – discontinuous signal peaks mapped in every other timeframe and found within a 400 µm Euclidean range were indirectly interconnected (Fig. 4C, Movie S11). The propagation trajectory between the discontinuous signal peaks was reconstructed using remnants of skeletonized signals. Signal amplitudes extracted from the corresponding electrical images were averaged over three consecutive timeframes (Δt = 150 µs) and mapped over interconnection areas. Obtained maps were next skeletonized to infer propagation trajectory between signal peaks found in the first and third timeframe. Conduction velocities estimated in the previous steps were used as criteria for selecting optimal propagation trajectories and predicting spatial coordinates of data for the second timeframe. Reconstruction of axonal trajectories over discontinuous signal peaks revealed axonal branching forks that could not be reconstructed in the previous steps.

### Performance of the method for reconstructing axonal functional morphology

The Bayes optimal template-matching technique was used to estimate the algorithm’s performance for detecting axonal AP peaks (Fig. 5). The technique can reliably discriminate axonal APs from the background noise^9^ and therefore provide a ground truth for the validation of the algorithm. Herein, template-matching was used to discriminate between ‘false peaks’ caused by the noise and ‘true peaks’ that resulted from axonal electrical activity (Fig. 5A). Discrimination criteria were based on similarities (‘matching’) between waveforms of averaged signals (‘templates’) and corresponding single trials. High similarities were expected in cases of accurate axonal signals (APs) and low similarities in cases where signals were derived from the background noise. We constructed templates for each signal peak in an electrical image. Constructed templates were next compared with waveforms of corresponding single trials, and a percentage of ‘matching’ trials was computed for each of the peaks. Signal peaks that yielded a match of > 70 % were classified as true peaks. Classified (true and false) peaks were used as the ground truth to estimate performances of the algorithm for detecting axonal AP peaks (Fig. 5B, C). Data obtained from 20 cells (ten cortical and ten motor neurons) were used for this analysis.

**Figure 5.**
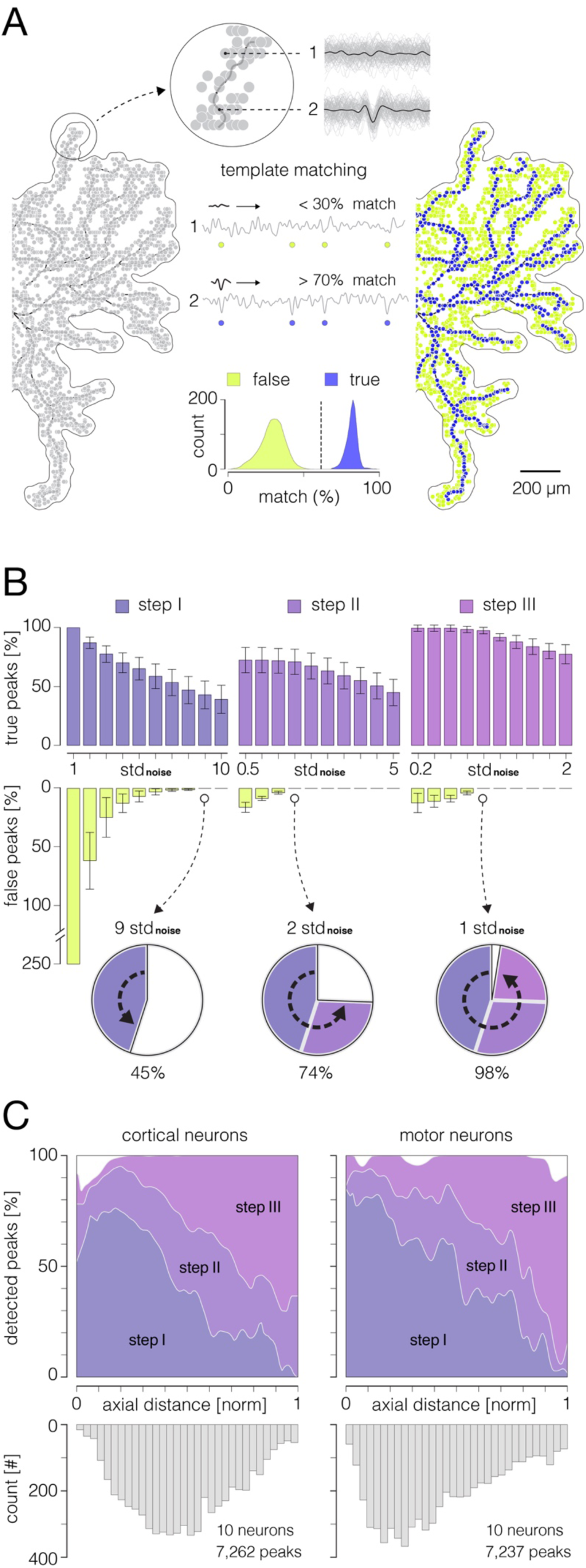
Performances of the algorithm for detection of axonal signal peaks. **(A)** Principle of the template matching demonstrated on data recorded from cortical axons: (Left) Extracellular APs were spike-sorted (see Fig. S6) and reconstructed across an entire array (see Fig. S7) by averaging 100-200 single-trials on each electrode. Reconstructed 3D signals were next transformed into derivatives (see Fig. 3) and all available signal peaks were obtained. Obtained peaks (pale-gray circles) are superimposed over the contour of axonal morphology. (Middle) Voltage traces recorded at locations of two example peaks are shown at the top (denoted as 1 and 2); averaged templates (black traces) are superimposed over single recording trials (light-gray traces). Note weak similarities in waveforms between the template and single trials at location 1 (putative noise) as oppose to strong similarities at location 2 (putative axonal signal). Template-matching procedure: the waveform of the template is successively compared with waveforms of discrete recording trials to extract percentage of “matching” trials within total number of trials. Examples of recording trials are presented continually within voltage traces (denoted as 1 and 2) and their relative positions are marked by circles: yellow and blue circles denote template mismatches and matches, respectively. Histogram below expresses separability between true and false peaks obtained from the template-matching; all obtained peaks (as shown in left) were used for this analysis. Data that yielded a match of > 70 % were considered true axonal signal. (Right) True and false peaks (blue and yellow circles, respectively) were classified using the template-matching technique. **(B)** Performance of the algorithm for AP peak detection. Histograms show percentages of true and false peak detections across the three thresholding steps (see Fig. 3) with varying threshold levels. Percentages of true peak detections are exposed through violet, purple and magenta ascending bars (step I, II and III, respectively); percentages of false peak detections are exposed through yellow descending bars. True and false peak detections are expressed as percentages of corresponding peaks detected by the template-matching. Note: no false peak detections were observed at thresholds set to 9, 2 and 1 STD of the noise in step I, II and III, respectively. Pie-charts below show progress in the true peak detection across the three steps. **(C)** Detectability of signal peaks across arbors of cortical and motor neuron axons. Detectability for the three steps is expressed over axial distances from the putative AISes. The bottom histogram shows spatial distribution of detected peaks. Data shown in (B, C) was obtained from 10 cortical and 10 motor neurons.

The performance of the algorithm for the detection of axonal AP peaks is shown in Fig. 5B. We estimated the performance of the three detection steps (see Fig. 3) with varying threshold levels. We were able to detect 45 %, 74 %, and 98 % of the actual peaks after the first, second and third steps, respectively. We observed no false peak detections for thresholds set to 9, 2, and 1 STD of the noise in 1^st^, 2^nd^ and 3^rd^ step, respectively. The detectability of AP peaks across axonal arbors of cortical and motor neurons is shown in Fig. 5C.

Axonal conduction trajectories, obtained from stimulation-triggered neuronal activity, were applied to estimate the performance of the algorithm for tracking axonal AP peaks (Fig. 6). The key concepts of the stimulation protocol used in this study are presented in Fig. S8 and described in the methods section. We used targeted stimulation to reveal and verify the same axońs conduction trajectory by altering its conductiońs direction (Fig. 6A, Movie S8). Stimulation-triggered APs were mapped spatially across all microelectrodes and temporally over discrete timeframes (with 50 µs inter-frame interval) to produce the movie. Observing the spatial movements of AP peaks in consecutive movie frames enabled us to track axonal conduction in different directions visually and to reconstruct axonal conduction trajectories (as shown in Fig. 6A and Movie S12, S13).

**Figure 6.**
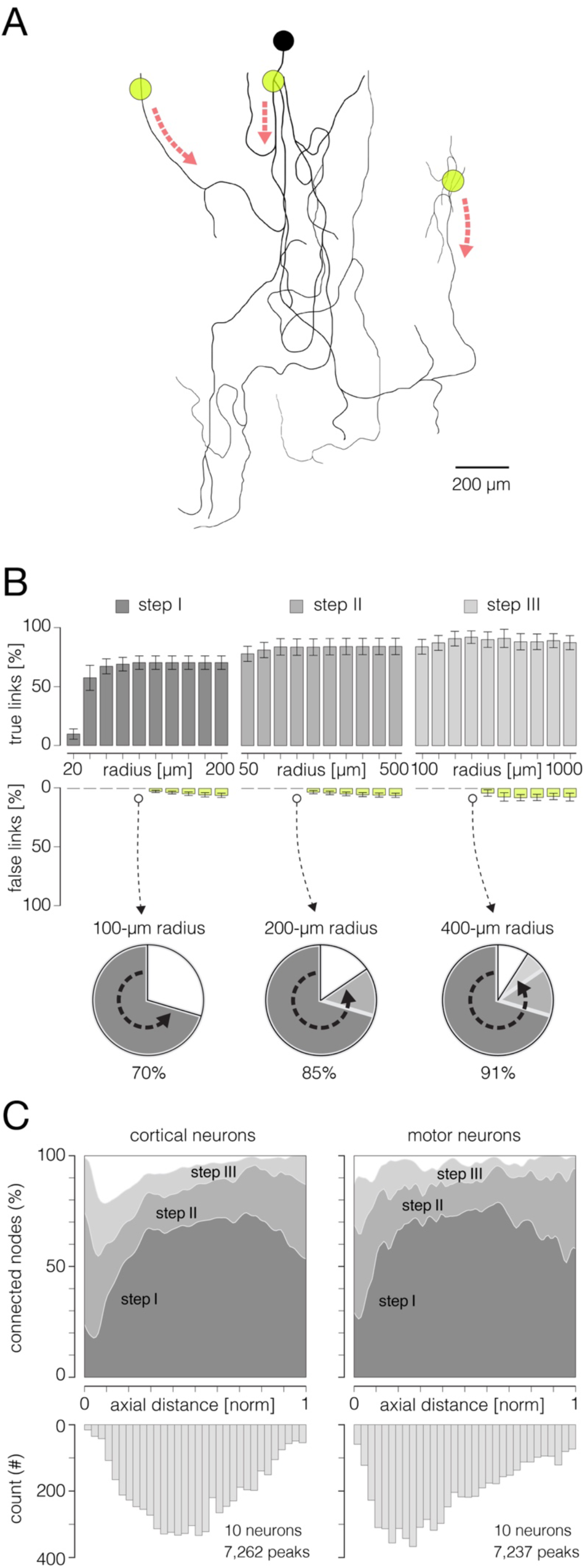
Performances of the algorithm for reconstruction of axonal functional morphology. **(A)** Axonal conduction trajectories (black axonal contour) reconstructed from stimulation-triggered neuronal activities; electrical stimulation targeted three different axonal sites (yellow semitransparent circles); red dashed arrows indicate directions of axonal conduction triggered from the corresponding stimulation site. Stimulation-triggered APs were reconstructed across an entire array and presented across consecutive movie frames, enabling visual tracking and reconstruction of axonal conduction trajectories (see Movie S12, S13). **(B)** Performance of the algorithm for reconstruction of axonal functional morphology. Histograms display percentages of true and false links established between mapped AP peaks using direct, skeletonization-assisted and indirect interconnection (see Fig. 3). Percentages of true links are exposed through dark-gray, gray and pale-gray ascending bars (step I, II and III, respectively); percentages of false links are exposed through yellow descending bars. Percentages of true and false links obtained using different radii for interconnection of the peaks are shown for each of the steps. The percentages are expressed with respect to a total number of links found in the corresponding ground truth trajectory (as shown in A). Note: no false links were established within Euclidean range (radius) of 100, 200 and 400 µm in step I, II and III, respectively. Pie-charts below show progress in the reconstruction of axonal morphology across the three steps. **(C)** Efficiency of the reconstruction (% of connected nodes) estimated for the three interconnection steps. Efficiencies are shown separately for cortical and motor neuron axons and are expressed over axial distances from the putative AISes. The bottom histogram shows spatial distribution of connected nodes. Data shown in (B, C) was obtained from 10 cortical and 10 motor neurons.

The algorithm’s performance for tracking axonal conduction is shown in Fig. 6B. We estimated the performance of the three tracking steps (see Fig. 4) with varying diameters of corresponding interconnection areas. We could interconnect 70 %, 85 %, and 91 % of the mapped AP peaks after the first, second and third step, respectively. We observed no false links when using the interconnection areas with diameters of 100, 200, and 400 µm in 1^st^, 2^nd^ and 3^rd^ step, respectively. The efficiency of the signal tracking across axonal arbors of cortical and motor neurons is shown in Fig. 6C.

### Functional morphologies of cortical and spinal axons

The reconstructed functional morphologies revealed axonal arbors of cortical and motor neurons and indicated positions of their AISes and somas (Fig. 7). They further provided insights into total axonal lengths, the spatial distribution of axonal branching-points and terminals, lengths of inter-branching segments and their branching orders. We used electrical images to extract areas occupied by axonal electrical activity (“active areas”).

**Figure 7.**
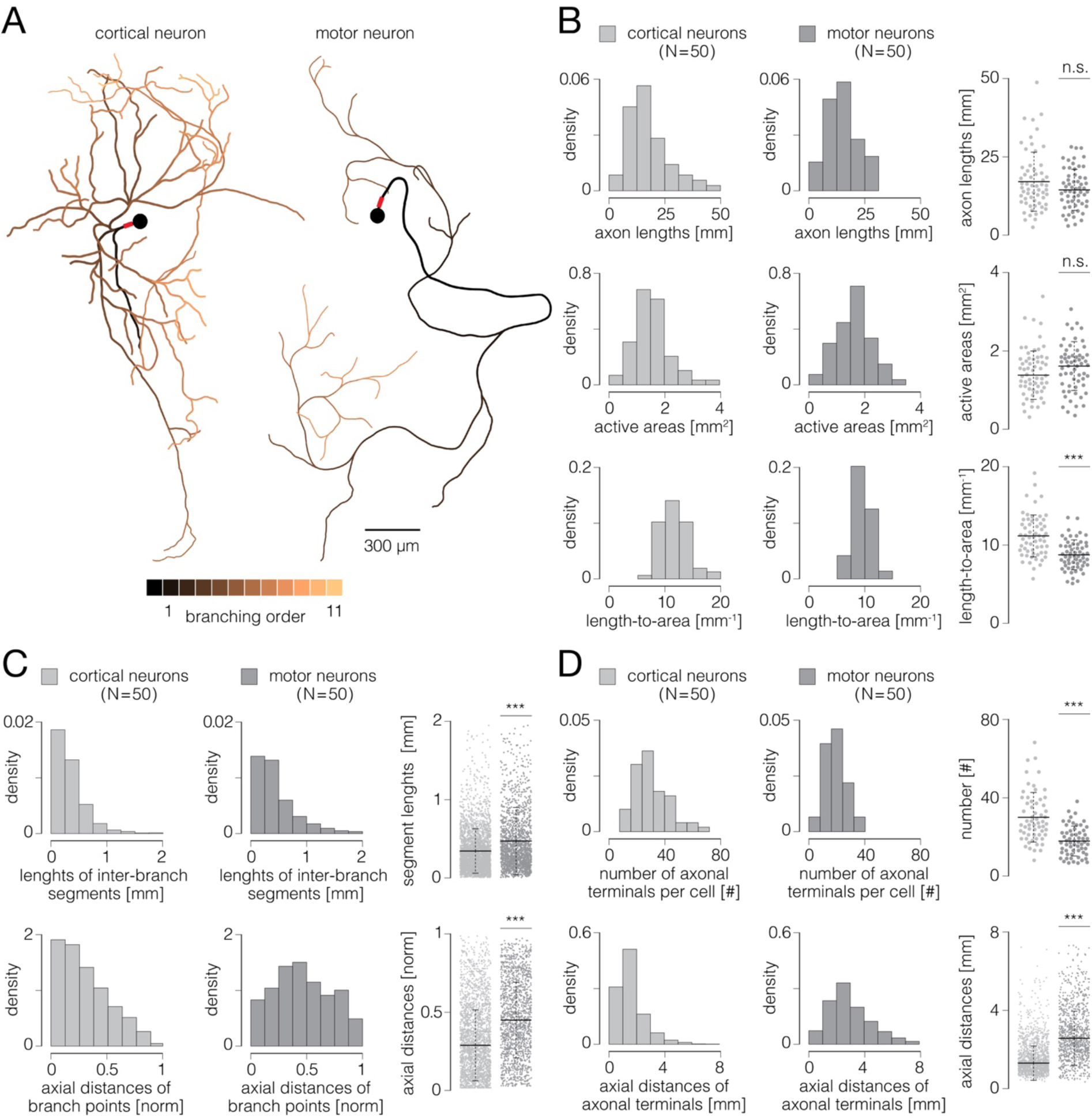
Functional morphologies of cortical and spinal axons. **(A)** Functional morphologies of a cortical (left) and motor-neuron axons (right) reconstructed based on spontaneous neuronal activities. Branching orders of reconstructed neurites are color-coded; neuronal somas are presented by black-filled circles. Axon initial segments are denoted by thick red lines near somas. Functional morphologies of the two neurons are also presented in Movie S14. **(B)** Axonal lengths and active areas. Histograms express density distributions of total axonal lengths, sizes of active areas and length-to-area quotients. Charts express comparisons between the corresponding values obtained from cortical and motor neurons; horizontal black lines denote mean values, perpendicular black-dashed lines denote standard deviations. **(C)** Lengths of inter-branching segments and axial distances of branching points. (Up) Histograms express density distributions of lengths of inter-branching segments. (Down) Histograms express density distributions of axial distances of branching points (with respect to corresponding AIS). Charts on the right express comparisons between the corresponding values obtained from cortical and motor neurons; horizontal black lines denote mean values, perpendicular black-dashed lines denote standard deviations. **(D)** Quantity and axial distances of axon terminals. (Up) Histograms express density distributions of numbers of axon terminals per cell. (Down) Histograms express density distributions of axial distances of axon terminals (with respect to corresponding AIS). Charts on the right express comparisons between the corresponding values obtained from cortical and motor neurons; horizontal black lines denote mean values, perpendicular black-dashed lines denote standard deviations. Color-code: pale-gray color was used to mark data obtained from cortical neurons; dark-gray color was used to mark data obtained from motor neurons. Data shown in (B, C, D) was extracted from functional morphologies of 50 cortical and 50 motor neurons. *** p < 0.001.

Representative examples of functional morphologies reconstructed for cortical and motor neurons are shown in Fig. 7A, and displayed in Movie S14. The reconstructed arbor of the cortical axon yielded a total length of 27.12 mm, comprising 101 inter-branching segments and 53 axon terminals. Axonal electrical activity was detected on 7,295 electrodes, occupying an active area of 2.23 mm^2^. The axonal arbor of the motor neuron yielded a total length of 15.26 mm, comprising 43 inter-branching segments and 23 axon terminals. Axonal activity was detected on 6,663 electrodes, occupying an active area of 2.04 mm^2^. We reconstructed functional morphologies for 50 cortical and 50 motor neurons and analyzed morphological features of their axonal arbors, a total length of 1.04 m and 0.81 m of cortical and spinal axons, respectively.

We found that cortical and motor neurons grew axons comparable in their total lengths. However, ratios between axonal lengths and their corresponding active areas were significantly higher for cortical than spinal axons (Fig. 7B). The average axonal lengths were 16.73±1.20 and 14.63±0.88 mm for cortical and motor neurons, respectively (p=0.25). The average sizes of active areas were 1.38±0.08 and 1.61±0.08 mm^2^ for cortical and motor neurons, respectively (p=0.23). The average length-to-area quotients were 11.72±0.32 and 9.49±0.24 mm^-1^ for cortical and motor neurons, respectively (p<10^-6^).

We found cortical axons to develop branching points in more proximal parts of their arbors and to form shorter inter-branching segments as compared to spinal axons (Fig. 7C). Average axial distances of axonal branching points were 0.27±0.01 and 0.43±0.01 mm for cortical and motor neurons, respectively (p<10^-6^). Average lengths of axonal inter-branching segments were 0.34±0.01 and 0.48±0.01 mm for cortical and motor neurons, respectively (p<10^-6^).

We found significantly more axon terminals in cortical than in spinal axons; however, spinal axons projected their terminals at significantly greater distances as compared to cortical axons (Fig. 7D). Average numbers of axon terminals were 29.92±1.64 and 17.82±1.03 for cortical and motor neurons, respectively (p<10^-6^). Average axial distances of axon terminals were 1.17±0.02 and 2.41±0.04 mm for cortical and motor neurons, respectively (p<10^-6^).

### Signal amplitude fluctuations during axonal conduction

In addition to exposing the axonal structure, functional morphologies also reveal APs as propagating across axonal arbors (Fig. 8). They allow observation of AP waveforms at any axonal location and mapping of AP amplitudes across entire arbors. Representative AP waveforms extracted from functional morphologies of cortical and spinal axons are shown in Fig. 8A, and displayed in Movie S15. We investigated fluctuations in AP amplitudes across functional morphologies of 50 cortical and 50 spinal axons (same neurons as in Fig. 7). The analysis included 45,232 data points obtained from 1.044 m of cortical axons and 36,286 data points obtained from 0.812 m of spinal axons.

**Figure 8.**
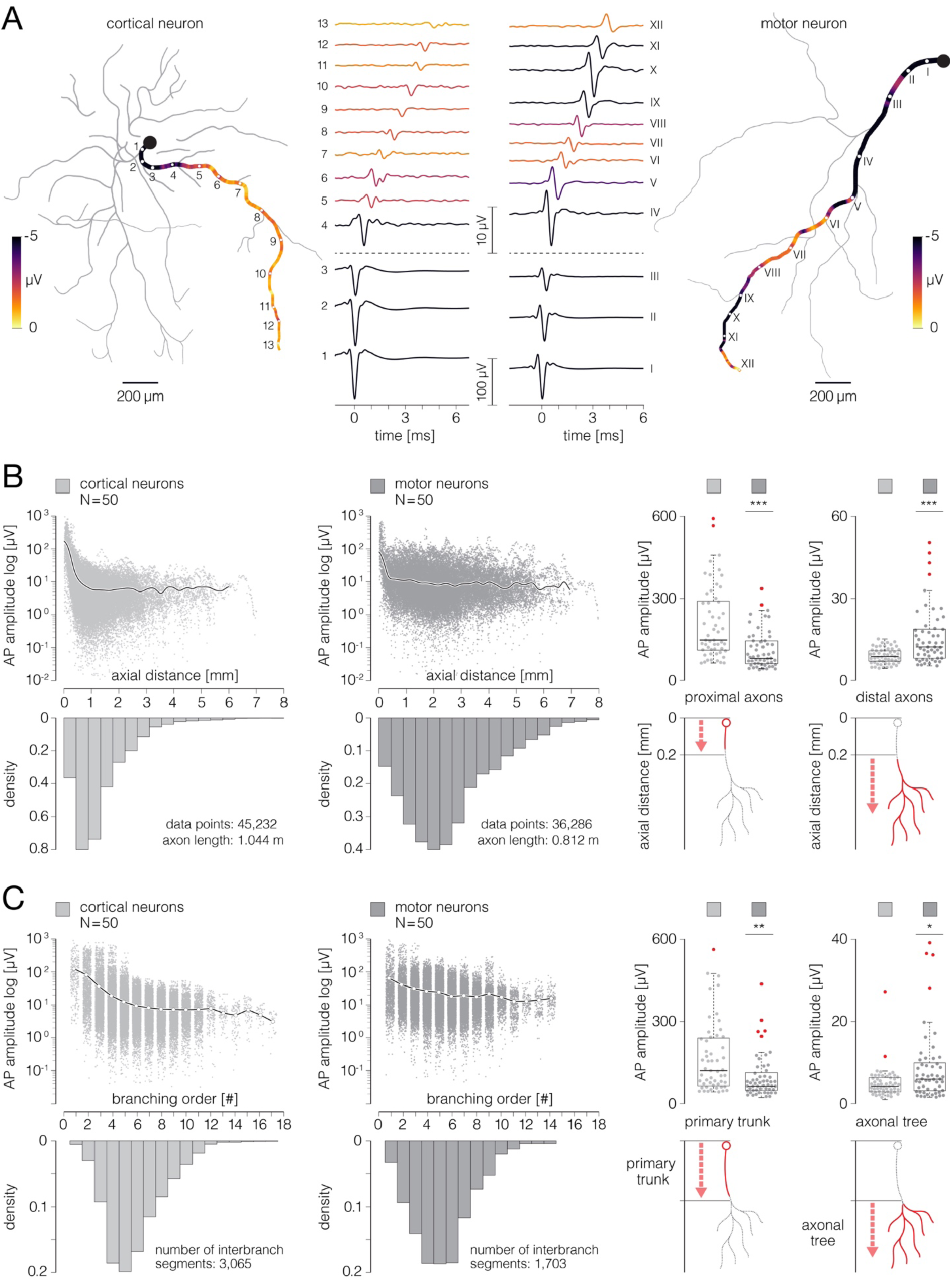
Signal amplitude fluctuations during axonal conduction. **(A)** Action potential waveforms extracted from functional morphologies of cortical and motor-neuron axons. (Left) Functional morphology of a single cortical neuron is displayed in gray; average AP amplitudes tracked across selected axonal path are color-coded; waveforms of axonal APs obtained from the denoted locations (numbered 1-13) are shown beside. (Right) Functional morphology of a single motor neuron is displayed in gray; average AP amplitudes tracked across selected axonal path are color-coded; waveforms of axonal APs obtained from the denoted locations (numbered I-XII) are shown beside. Note difference in scalebars for AP waveforms obtained from proximal and distal axons. Data from the two neurons are also presented in Movie S15. **(B)** Axonal AP amplitudes versus axial distance from the AIS. (Left-middle) Amplitudes of axonal APs versus axial distances of their recording sites are plotted for cortical and motor neurons; axial distances are expressed with respect to locations of the corresponding AISes. The black curves represent mean values of amplitudes over all data points. The bottom histograms show density distributions of axial distances from the corresponding AISes. (Right) Mean AP amplitudes obtained from proximal and distal axons are compared between cortical and motor neurons; comparisons are expressed using box plots. The bottom diagrams show criterium for discriminating proximal from distal axonal locations. Action potential amplitudes averaged over proximal and distal regions of each of the neurons were used in this analysis. **(C)** Axonal AP amplitudes versus axonal branching order. (Left-middle) Amplitudes of axonal AP versus branching order of corresponding axonal segments are plotted for cortical and motor neurons. The white circles (interconnected by black lines) represent mean values of amplitudes computed for discrete branching orders. The bottom histograms show density distributions of branching orders. (Right) Mean signal amplitudes obtained from primary axonal trunks and axonal trees are compared between cortical and motor neurons; comparisons are expressed using box plots. The bottom diagrams show criterium for discriminating primary axonal trunks from axonal trees. Color-code: pale-gray color was used to mark data obtained from cortical neurons; dark-gray color was used to mark data obtained from motor neurons. Data shown in (B, C) was extracted from functional morphologies of 50 cortical and 50 motor neurons. ** p < 0.01; *** p < 0.001.

We found that APs recorded from the most proximal parts of axons had much higher amplitudes than APs recorded from more distal axonal locations (Fig. 8A-C). To further compare AP amplitudes across different cortical and spinal axon domains, we segregated the reconstructed morphologies into proximal and distal axons, primary trunks, and axonal trees. Proximal axons pertain to locations found within the first 0.2 mm of axial length; all other locations (axial distance > 0.2 mm) were considered as distal (see Fig. 8B-right). Primary trunk entails locations found in the axonal domain between the soma and the first branching fork, all other locations were assigned to the axonal tree (see Fig. 8C-right).

APs recorded from proximal axons had significantly higher amplitude in cortical (44.58±4.81 µV) than in motor neurons (19.07±2.53 µV; p<10^-6^). On the contrary, APs recorded from distal axons had significantly smaller amplitude in cortical (1.50±0.10 µV) than in motor neurons (3.30±0.38 µV; p<10^-6^; Fig. 8b).

Similarly, APs obtained from primary axonal trunks had significantly higher amplitude in cortical than motor neurons, whereas, APs obtained from axonal trees had significantly lower amplitude in cortical than motor neurons (Fig. 8C). Average amplitudes of APs recorded from primary axonal trunks were 33.29±4.78 and 15.06±2.78 µV for cortical and motor neurons, respectively (p=0.002). Average amplitudes of APs recorded from axonal trees were 1.25±0.13 and 2.17±0.30 µV for cortical and motor neurons, respectively (p=0.024).

### Temporal dynamics of axonal conduction

Functional morphologies carry information about times at which APs arrived at any axonal site and, as such, provide direct insights into temporal dynamics of axonal conduction (Fig. 9). Axonal functional morphology reveals time at which axonal conduction is initiated (“initial time”) and allows to obtain times at which axonal APs arrive at axon terminals (“arrival time”). It enables extraction of the interval between the earliest and the latest arrival time (“interval of arrivals”) and inspection of the total duration of the axonal conduction (“active timespan”) (see Fig. 9A-left). Functional morphologies allow for the inspection of AP conduction dynamics across entire axonal arbors and over individual axonal paths (see Fig. 9A-right). Finally, structural and temporal parameters from functional morphologies enable computation of conduction velocities for any axonal segment.

**Figure 9.**
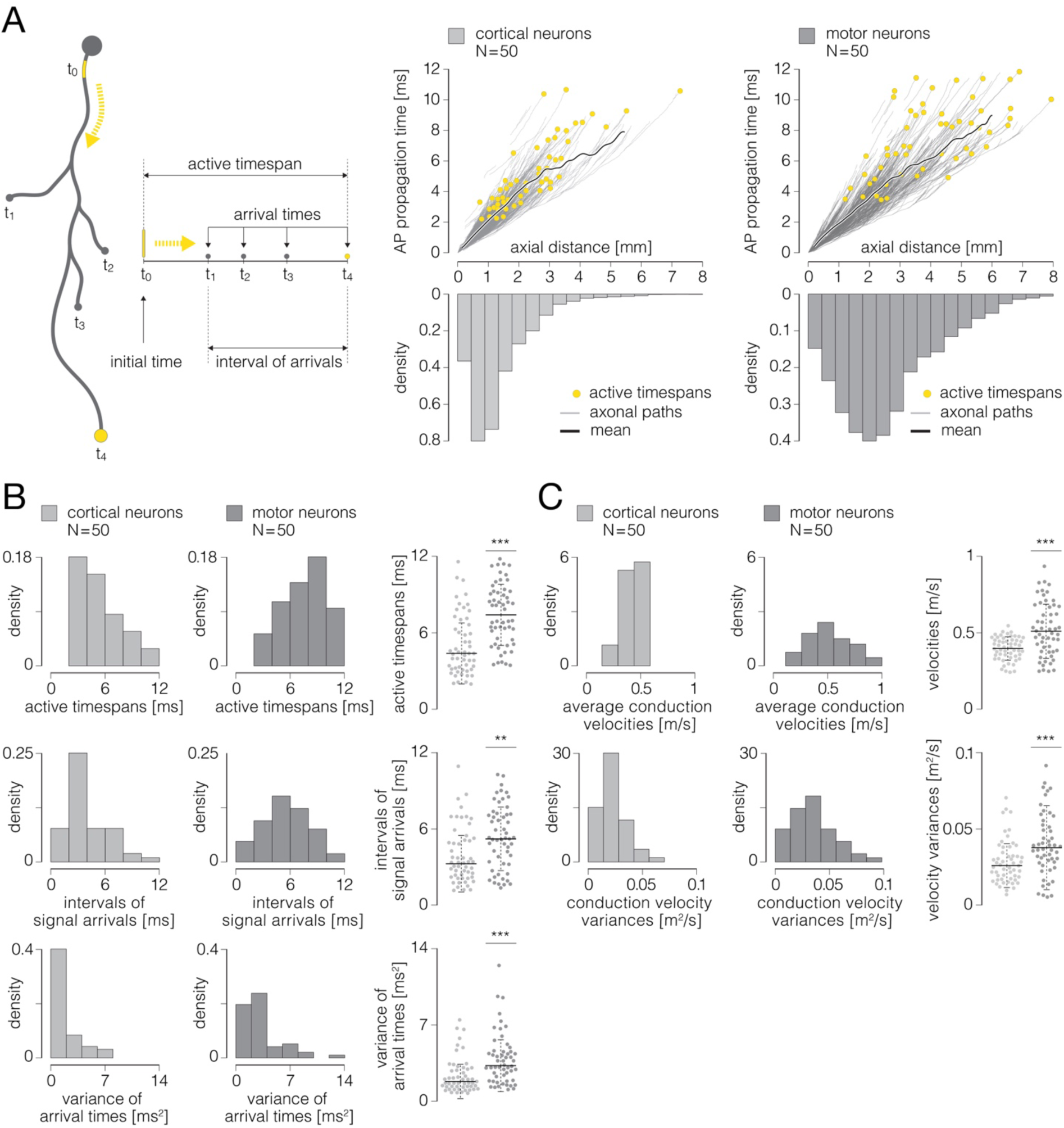
Temporal dynamics of axonal conduction. **(A)** (Left) Graphical representation of parameters that were used for the analysis of temporal aspects of axonal conduction. Thick gray contour represents simplified axonal morphology; neuronal somas are presented by gray-filled circles; a yellow line superimposed over a proximal axon represents the AIS; blebs at the ends of four axonal branches represent axon terminals. Yellow-dashed arrows indicate directionality of axonal AP propagation. Temporal parameters: initial time (t_0_) - time of the initiation of axonal conduction; arrival times (t_1-4_) - times at which axonal APs arrive at axon terminals; interval of arrivals - interval between the earliest and the latest arrival time; active timespan - timespan between initial time and the latest arrival time. (Right) Axonal AP propagation time versus axial distance from the AIS - comparison between cortical (graph on the left) and motor neurons (graph on the right). Yellow circles represent active timespans; gray curves in the background represent AP propagation times observed over individual axonal paths; thick black curves represent mean values of propagation times averaged over all axonal paths. **(B)** Temporal aspects of axonal conduction. (Up) Histograms express density distributions of active timespans for cortical and motor neurons. (Middle) Histograms express density distributions of intervals of arrivals for cortical and motor neurons. (Down) Histograms express density distributions of variances of arrival times for cortical and motor neurons. Charts on the right express comparisons between the corresponding values obtained from cortical and motor neurons; horizontal black lines denote mean values, perpendicular black-dashed lines denote standard deviations. **(C)** Axonal conduction velocities. (Up) Histograms express density distributions of average conduction velocities for cortical and motor neurons; average velocities are computed for individual cells. (Down) Histograms express density distributions of variances of conduction velocities for cortical and motor neurons; variances are computed for individual cells. Charts on the right express comparisons between the corresponding values obtained from cortical and motor neurons; horizontal black lines denote mean values, perpendicular black-dashed lines denote standard deviations. Color-code: pale-gray color was used to mark data obtained from cortical neurons; dark-gray color was used to mark data obtained from motor neurons. Data shown was extracted from functional morphologies of 50 cortical and 50 motor neurons. ** p < 0.01; ***p < 0.001.

We investigated temporal aspects of axonal conduction across functional morphologies of 50 cortical and 50 spinal axons (same neurons as in Fig. 7 and Fig. 8). The analysis included 45,232 data points obtained from 1.044 m of cortical axons and 36,286 data points obtained from 0.812 m of spinal axons.

Cortical neurons had significantly shorter active timespans and higher synchronization of the arrival times as compared to motor neurons (Fig. 9B). Active timespans were 4.60±0.33 and 7.40±0.32 ms for cortical and motor neurons, respectively (p<10^-6^). Intervals of signal arrivals at axon terminals were 3.44±0.31 and 5.40±0.34 ms for cortical and motor neurons, respectively (p=0.0024). Variance of signal arrival times were 0.95±0.23 and 2.50±0.35 ms^2^ for cortical and motor neurons, respectively (p<10^-4^).

We next computed axonal conduction velocities using structural and temporal parameters obtained from the functional morphologies. Conduction velocities were calculated from axial distances versus AP propagation times (Fig. 9A-right), and were computed over 100 µm long axonal chunks stepped by 17.5 µm.

We found cortical axons to exhibit significantly slower and more uniform conduction velocities compared to spinal axons (Fig. 9C). Average conduction velocities were 0.46±0.01 and 0.56±0.02 m/s for cortical and motor neurons, respectively (p<10^-4^). Variances of conduction velocities were 0.03±0.002 and 0.04±0.003 m/s for cortical and motor neurons, respectively (p<10^-4^).

## DISCUSSION

We developed a method for noninvasive functional imaging of unmyelinated mammalian axons *in vitro* using the CMOS-based HD-MEA system (Fig. 1, Movie S1, S2). The method yields an axonal ‘functional morphology’, comprising multidimensional data derived from extracellular APs recorded during axonal conduction. Functional morphology contains information about axonal conduction trajectories mapped at high spatial and temporal resolution (Movie S6-11). It allows the reconstruction of axonal ‘electrical morphology’ at different timepoints during the conduction and, at the same time, to expose waveforms of the propagating APs (Fig. 8, Movie S15).

The presented method has been developed and validated on primary rodent cortical and motor neurons cultured directly over HD-MEA surfaces (Fig. S1-3). The cultures matured within two weeks and exhibited spontaneous electrical activities (Fig. S6). They efficiently adhered to HD-MEA surfaces and formed monolayers that varied in local thicknesses and cell-densities (Fig. S4). Neurons developed complex axonal arbors and provided a tight interface between axons and microelectrodes. Because axons grew in a 3D manner, their distances from the HD-MEA surfaces varied locally (Fig. S5), but remained within the sensing range of the microelectrodes^32^.

Spontaneous electrical activity enabled us to map individual neurons in the cultures (Fig. S6) and expose the spatiotemporal distribution of APs recorded across their axonal arbors (Fig. S7, Movie S5). Mapping individual neurons was possible thanks to high-amplitude APs recorded at their AISes ^29, 30^. The AIS signals broadly surpass the background noise and hence could be easily detected^9^. Because they colocalized with the AIS, local maxima found within activity maps (Fig. S6A) indicate proximal regions of individual neurons in the network^29, 30^. Owing to the high-density arrangement of recording microelectrodes (Fig. S1)^33^, spike-sorting enabled discerning mixed signals recorded from neighboring neurons (Fig. S6B) and thereby extracting APs arising from individual neurons (Fig. S6C). Averaging of extracted signals over all electrodes revealed the spatiotemporal distribution of a single neuron’s activity. Such signal representation was referred to as an “axonal electrical image” (Fig. S7, Movie S5) and it carries information about neuronal subcellular elements^29, 30^. Thus, the largest and the earliest signals in the footprint arise from the AIS, much smaller and fast propagating signals arise from axons, and slow back-propagating signals arise from the soma and proximal dendrites^29, 30^. Thanks to the low-noise CMOS-design, the HD-MEA chip allows detection of AP propagation across large portions of entire axonal arbors^9^. The soma and proximal dendrites provide minor contributions to the electrical image, typically masked by larger AIS signals^29^. Owing to their low-amplitude extracellular signals, distal dendrites do not seem to be detectable with the HD-MEA system used here.

Adaptive thresholding applied to axonal electrical images enabled us to map extracellular APs propagating across axonal arbors (Fig. 3, Movie S6-8). Signal peaks were traced at high spatial and temporal resolution thanks to the dense arrangement of the electrodes and high-frequency sampling rate^33^. The thresholding scheme was constructed to detect signals with various amplitudes while minimizing detection errors. Simple planar thresholds, set to 9 STD of the background noise, were used to detect high-amplitude signal peaks (Fig. 3A, Movie S6). Since these thresholds provided detection high above the level of the background noise, they could be applied across an entire array while yielding no detection errors. They, however, failed to detect low-amplitude signal peaks and only provided fragmentary detection. Instead, spatiotemporal confinement of low-level thresholds, contingent on coordinates of high-amplitude peaks, enabled the mapping of low-amplitude signals (Fig. 3B, Movie S7). This was possible thanks to the very nature of axonal conduction, which can be represented using a simplified Markov chain model^34^. Namely, because signal propagation in unmyelinated axons is continuous, it represents a cascade of successive events in which the probability of each event depends solely on the state attained in the preceding event. Therefore, there was a high probability of detecting signal peaks in the immediate spatiotemporal proximity of previously mapped peaks. While confining the reach of the thresholds to local areas minimized the influence of the background noise, temporal confinement to preceding and following timeframes excluded signals that occurred much earlier and later than the reference peak.

Further relaxation of the threshold confinements enhanced the detection of low-amplitude signals but inevitably increased the risk of detection errors (Fig. 3C, Movie S8). In principle, our adaptive thresholding scheme followed a greedy algorithm paradigm^35^, and as such, it is vulnerable to detection errors. Generally, a greedy algorithm builds up a global solution stepwise by making the locally optimal choice at each step. However, erroneous solutions made at earlier steps can proliferate and thereby compromise the global solution^36^. Therefore, carefully tuned thresholding parameters are key to the optimal detection of signal peaks.

The approach of multiple-moving object tracking enabled us to reconstruct axonal conduction trajectories through iterative associations of the mapped signal peaks (Fig. 4, Movie S9-11). The tracking algorithm was designed to estimate inter-frame correspondences between the mapped peaks in a data-driven fashion. It operates within predefined spatiotemporal boundaries, but also directly learns data-association criteria from sequences reconstructed along the tracking process. Thanks to strict boundaries, the algorithm discarded trajectories yielding conduction velocities that are unlikely for mammalian axons. While such boundaries ensure the exclusion of slow-propagating dendrites^29^, they could also omit super-fast saltatory conduction inherent to myelinated axons^10^, which were most likely not present in our preparations. Owing to the data-driven refinement of the association criteria, the algorithm allowed customization of the tracking process for a specific neuron. Such customization provides applicability of the algorithm to axons with different conduction velocities, allowing observation of variations between neurons of the same or different types.

Considering the Markov model of axonal conduction, AP peaks mapped in immediate spatiotemporal proximities can be associated directly using a greedy nearest neighbor matching (Fig. 4A, Movie S9). Due to the limited reach of the matching caliper, direct interconnection enabled only reconstruction of short fragments of the axonal conduction trajectory. Reconstructed fragments, in return, enabled estimation of the conduction velocity and thereby refining the association criteria. Because conduction velocities vary across a single axon^9,11^, greedy matching was limited to peaks mapped closely over sections of the trajectory with relatively constant conduction velocities. Relaxation of the association criteria could potentially extend the reach of the matching. However, it would inevitably introduce erroneous interconnections between spatially segregated peaks stretched over notably faster sections of the trajectory.

Skeletonization of an axonal electrical image enabled reproduction of conduction paths between distant peaks (Fig. 4B, Movie S10). Because skeletal remnants reflect the spatiotemporal landscape of the axonal signals, they can optimize the association of mapped peaks graphically, thus sidestepping complex heuristic techniques needed for maintaining computational tractability. In general, topology and structural complexity of skeletal remnants depend on spatiotemporal boundaries within which the axonal signals were skeletonized. Narrow boundaries encompassing two consecutive timeframes captured brief sequences of the axonal conduction and yielded skeletal remnants with fairly simple structure. Simple remnants enabled interconnection of the peaks across consecutive timeframes, however, discontinuous peaks remained beyond their reach.

Skeletonization within wider spatiotemporal boundaries allowed interconnecting peaks across discontinuous timeframes (Fig. 4C, Movie S11), but yielded more complex remnants, especially in regions where axons branched or intertwined. Complex remnants provided multiple possible solutions for the peak associations, thus aggravating the selection of an optimal conduction path between corresponding peaks. By restricting the skeletonization to three consecutive timeframes and using the refined association criteria, we could select the optimal path in most cases. However, further widening of the skeletonization boundaries increased the number of possible solutions exponentially and inevitably led to a deterioration of tracking performance.

The Bayes optimal template-matching technique was used to validate the algorithm for signal peak detection (Fig. 5). We chose this technique because it has previously been demonstrated to provide reliable detection of extracellular APs recorded using the HD-MEA system^9^. It enables the detection of axonal APs within single recording trials and microsecond differences in axonal propagation^9, 37^. Bayes optimal template-matching is sensitive enough to discriminate low-amplitude APs from the background noise^9^, and thus can be used to validate our algorithm.

Stimulation-triggered axonal activities allowed us to trace axonal conduction trajectories accurately and thus provided the ground truth for validation of our tracking algorithm (Fig. 6, Movie S12, S13). We have previously shown that both spontaneous and stimulation-triggered neuronal activities recorded by HD-MEAs can be used to outline axonal conduction trajectories^27, 30^. Moreover, we demonstrated a congruence between the reconstructed trajectories and actual axonal morphology revealed optically^9, 11, 27, 30^.

We investigated functional morphologies of cortical and spinal axons and found significant differences between their structures, AP amplitudes and conduction dynamics (Fig. 7-9), which potentially reflect biological hallmarks of different neuronal subtypes. The structural complexity of cortical axons was reflected in their extensive branching, followed by the abundance of the axon terminals that were projected locally and distally (Fig. 7, Movie S14). Such morphological features of cortical axons can be attributed to their role in providing synaptic connectivity within local neuronal assemblies as well as across large neuronal networks, thus enabling higher-order operations performed in the brain^38^. Spinal motor neurons, however, have evolved much simpler morphology to provide fundamentally different biological roles. Motor neurons innervate and precisely control muscles in the periphery outside the central nervous system and are, for that purpose, equipped with the longest known axons in the body^1^. Our results obtained in *in vitro* conditions are consistent with such a notion. They indicate that spinal axons indeed have much simpler morphologies when compared to cortical axons, yet still retain relatively long total lengths (Fig. 7, Movie S14). We found that spinal axons had moderately branched arbors, but projected their axon terminals at great distances, thus resembling morphological adaptations observed *in vivo*. Thanks to their extensively branched axons, cortical neurons enabled the delivery of APs to numerous axon terminals, surprisingly synchronously and within short timespans. On the contrary, spinal axons with less branches at their disposal required considerably more time to forward their APs to axon terminals in a less synchronized fashion (Fig. 9B).

We found that extracellular APs recorded from proximal axons near the putative AIS exposed disproportionately larger amplitudes than signals obtained from other axonal parts (Fig. 8, Movie S15). It has been previously shown that the AIS is the dominant contributor to the neuron’s extracellular electrical landscape^29^, which can be attributed to the high densities of voltage-gated ion channels expressed in the AIS ^39–41^. Interestingly, we found that APs obtained from proximal axons had significantly higher amplitudes in cortical than in motor neurons (Fig. 8B, C). The relatively low AP amplitudes at the AIS in motor neurons could result from the potentially disrupted organization of the AIS in unmyelinated motor neurons. Such disruption pertains to altered distribution and combination of voltage-gated ion channels as well as incomplete segregation of the AIS, para-AIS, and juxtapara-AIS compartments inherent to myelinated spinal axons^42^. We found that signals obtained from distal axons and axonal trees had significantly lower amplitudes in cortical than motor neurons (Fig. 8B, C). This difference could be explained by the fact that spinal axons have a considerably larger diameter than cortical axons^13^. As such, they engage a greater number of voltage-gated ion channels and create stronger transmembrane currents during an AP electrogenesis^43^.

Large diameters of spinal axons could also explain their significantly faster conduction velocities compared to cortical axons (Fig. 9C). In general, axonal diameter and the presence of myelin sheaths are crucial factors that control conduction velocity in mammalian axons^12, 44^. On the one hand, conduction velocity of unmyelinated axons is proportional to the square root of the axon diameter^2^. On the other hand, myelination provides a fundamentally different mechanism of AP propagation known as “saltatory” conduction^45^. In this case, myelin sheaths spatially restrict the distribution of voltage-gated ion channels to nodes of Ranvier and thus impose discontinuous AP propagation up to 100-fold faster than propagation in unmyelinated axons^12, 13^. Culturing protocols used in this study do not anticipate the growth of Schwann cells needed for formation of myelin sheaths. Therefore, it is unlikely that complete myelination occurred in our cultures. We, however, cannot exclude potential cases where axons were partially myelinated. Moreover, incomplete myelinization or discontinuous distribution of voltage-gated ion channels could explain relatively large variances of conduction velocities in spinal axons (Fig. 9C).

Generally, standard deviations of the data presented in Fig. 7-9 could result from morphological and biophysical differences between various neuronal subtypes that may have existed in our cultures. Thus, for example, developed cortical cultures comprise excitatory glutamatergic and inhibitory GABAergic neurons which are known to differ in size and morphological complexities^46^. Even greater diversity of neuronal subtypes can be found among motor neurons, including alpha, beta, and gamma motor neurons that are structurally specialized to innervate different muscle fiber types in the body^1^. However, another question is to what extend such specializations can be recapitulated *in vitro* and whether electrophysiological parameters observed in cultures allow for discriminating between neuronal subtypes.

The presented method for functional imaging of cortical and spinal axons provides direct insights into the biophysical properties of axonal conduction. It allows investigation of interdependences between axonal function and structure, noninvasively and over extended periods. Functional morphologies contain data obtained from large portions of axonal arbors and, as such, allow studying the fidelity of signal conduction in different axonal elements, including conduction failures at branching points^10^ and signal attenuation in tiny axon terminals^14^. They also allow the inspection of activity-dependent modulation of extracellular AP waveforms^9, 47^ and analysis of consequent changes in times at which modulated signals arrive at axon terminals^9^. Owing to its noninvasive nature, the method enables the study of structural and functional changes in axons over long periods. Thus, for example, comparative analysis of functional morphologies obtained at different timepoints allows investigation of biophysical changes during axonal outgrowth and pathfinding - developmental processes that guide axons toward specific targets and enable wiring within neuronal networks^48, 49^. Complemented with techniques for electrical microstimulation (see Fig. S8), the method can be used to study activity-dependent axonal plasticity, including structural alteration such as the positional shift of the AIS^50, 51^ as well as functional adaptations that involve changes in axonal conduction velocities^11^. Because the HD-MEA system allows observing electrical activities across entire neuronal networks (see Fig. S6), the method can serve as a complementary tool for studying the functional interplay between network dynamics and plastic adaptations in axons^50, 52^.

In summary, the presented method is designed for specific tissue-on-a-chip platforms and may serve as an attractive tool for pharmacological and preclinical studies of neurodegenerative diseases^53^. The key potential of the method lies in its ability to gain multilevel knowledge of neuronal functionality, which may serve as a valuable asset in drug discovery and safety pharmacology. The method holds great potential for developing disease-specific bioassays, especially considering the advent of human induced pluripotent stem cell (iPSC) technology that allows for utilizing patient-derived neurons.

## METHODS

### Animal use

All experimental protocols were approved by the Uppsala Animal Ethical Committee under animal license C97/15 and follow the guidelines of the Swedish Legislation on Animal Experimentation (Animal Welfare Act SFS 2009:303) and the European Communities Council Directive (2010/63/EU).

### HD-MEA and signal processing

A complementary-metal-oxide-semiconductor (CMOS)-based HD-MEA system (MaxWell Biosystems AG) was used for extracellular neuronal recording and stimulation (Fig. S1). The array comprises 26,400 platinum microelectrodes (9.3×5.45 µm^2^) packed within an area of 3.85×2.10 mm^2^, providing a density of 3150 electrodes per mm^2^ (17.5 µm center-to-center pitch). Owing to a flexible switch matrix technology, up to 1024 readout and/or stimulation channels could be routed to the desired electrodes and reconfigured within a few milliseconds. On-chip circuitry was used to amplify (0-80 dB programmable gain), filter (high pass: 0.3-100 Hz, low pass: 3.5-14 kHz), and digitize (8-bit, 20 kHz) the neuronal signals. Digitized signals were sent to a field-programmable gate array (FPGA) board and further streamed to a host PC for real-time visualization and data storage. Recorded signals were up-sampled to 200 kHz following the Whitaker-Shannon interpolation formula. Python 3.7 and Matlab R2020a were used for data analysis and to design experimental protocols.

### Cortical cultures

We used a culturing protocol for the long-term maintenance of neural cultures^54^ (Fig. S2). Minor adaptations were introduced to the protocol to constrain culture growth to the sensing area of the array and maintain optimal growth media conditions during long-term experimentation^30^. Cortices from embryonic day 18 Sprague Dawley rat were dissociated enzymatically in trypsin supplemented with 0.25% EDTA (Thermo Fisher Scientific) and physically by trituration. The remaining cell aggregates and debris were filtered using 40-µm cell strainer (Corning). For cell adhesion, a layer of 0.05% polyethyleneimine (Sigma-Aldrich) in borate buffer (Chemie Brunschwig), followed by a layer of 0.02 mg ml^-1^ human recombinant laminin (BioLamina) in DPBS containing Ca^2+^ and mg^2+^ (Thermo Fisher Scientific) was deposited on the electrode array. To constrain culture growth to the electrode array, a cell-drop containing ∼50,000 cells and covering ∼4 mm^2^ was seeded in the center of the array. The plating media were changed to growth media after 24 h and regularly changed every 6 days. Plating media consisted of Neurobasal, supplemented with 10% horse serum (HyClone), 0.5 mM GlutaMAX and 2% B27 (Thermo Fisher Scientific). The growth media consisted of DMEM, supplemented with 10% horse serum (HyClone), 0.5 mM GlutaMAX and 1 mM sodium pyruvate (Thermo Fisher Scientific). Cultures were maintained inside an incubator under controlled environmental conditions (36°C and 5% CO_2_). The culturing chambers were sealed with a ∼1 mm layer of light mineral oil (Sigma-Aldrich) floating above the growth medium. The sealing provided selective permeability to gases, such as O_2_ and CO_2_, and prevented evaporation and consequent changes in the growth media’s osmolarity during long-term experiments.

### Motor neuron cultures

Protocol for growing motor-neuron cultures on HD-MEA surface is illustrated in Fig. S3. Spinal cords isolated from embryonic day 14 Sprague Dawley rat were dissociated enzymatically in trypsin supplemented with 0.25% EDTA (Thermo Fisher Scientific) and physically by trituration. The remaining cell aggregates and debris were filtered out using 40-µm cell strainer (Corning). A density gradient medium composed of 15% OptiPrep^TM^ (Sigma-Aldrich) in Leibovitz’s L-15 medium (Thermo Fisher Scientific) was used to fractionate motor neurons. For cell adhesion, a layer of 0.05% polyethyleneimine (Sigma-Aldrich) in borate buffer (Chemie Brunschwig), followed by a layer of 0.02 mg ml^-1^ human recombinant laminin (BioLamina) in DPBS containing Ca^2+^ & mg^2+^ (Thermo Fisher Scientific) was deposited on the electrode array. To constrain culture growth to the electrode array, a cell-drop containing ∼50,000 cells and covering ∼4 mm^2^ was seeded in the center of the array. The growth media consisted of Neurobasal supplemented with 20% MyoTonic differentiation medium (Cook MyoSite), 1% fetal bovine serum, 2% horse serum, 2% B27, 1% Antibiotic-Antimycotic, 0.2 mM Gentamicin, 0.7 mM L-glutamine, 1.5 mM sodium pyruvate (Thermo Fisher Scientific), 0.8 nM brain-derived neurotrophic factor (BDNF), 0.2 nM glial cell line-derived growth factor (GDNF), 0.2 nM ciliary neurotrophic factor (CNTF), 1.5 nM neurotrophin 3 (NT3), 1.4 nM neurotrophin 4 (NT4, R&D systems) and 1.3 nM human insulin-like growth factor 1 (hIGF-1, Thermo Fisher Scientific). The growth media were changed after 24 h and further regularly changed every 4 days. Cultures were maintained inside an incubator under controlled environmental conditions (36°C and 5% CO_2_). The culturing chambers were sealed with a ∼1 mm layer of light mineral oil (Sigma-Aldrich) floating above the growth medium.

### Immunocytochemistry

Neuronal cultures were fixed in 4% paraformaldehyde (Thermo Fisher Scientific) in PBS (Sigma-Aldrich) at pH 7.4 for 15 min at room temperature, washed twice with ice-cold PBS, permeabilized with 0.25% Triton X-100 (Sigma-Aldrich) in PBS for 10 min and washed three times in PBS. Fixed cultures were exposed to phosphate-buffered saline with Tween 20 (1% bovine serum albumin and 0.1% Tween 20 in PBS; Sigma-Aldrich) for 30 min to prevent unspecific binding of antibodies. The primary antibodies Anti-Map2 (Abcam, ab5392), Anti-β-III tubulin (Abcam, ab18207) and Anti-mCherry (Abcam, ab125096), diluted in phosphate buffered saline with tween 20 to ratios of 1:1000, 1:500 and 1:500 respectively, were added and left overnight at 4°C on a shaker. Cultures were washed three times in PBS for 5 min each time on the shaker. The secondary antibodies Alexa Fluor 488 (Abcam, ab150173), Alexa Fluor 488 (Abcam, ab150073) and Alexa Fluor 594 (Abcam, ab150116) diluted to ratio of 1:200 in PBS with 1% BSA were added and left for 60 minutes in the dark at room temperature. Samples were washed three times in PBS for 5 min in the dark and then stored at 4°C. Cultures were immunostained with anti-β-III tubulin (Fig. S4, S5 and Movie S1, S2), anti-Map2 (Fig. S6, S7 and Movie S3-5) and a combination of anti-mCherry and anti-β-III tubulin antibodies (Fig. S5).

### Live imaging

Live-cell visualization of neurons was performed by transfection using pAAV-hSyn-mCherry plasmid from Karl Deisseroth (Addgene plasmid # 114472) and Lipofectamine 3000 (Thermo Fisher Scientific), jetPRIME (Polyplus) or TurboFect (Thermo Fisher Scientific) transfection reagent following the manufacturer’s protocols. Micrographs of cortical and motor neurons transfected using Lipofectamine 3000 are shown in Fig. S5.

### Microscopy and 3D image reconstruction

A Nikon Eclipse LVDIA-N microscope, Nikon DS-Fi2 camera and the Nikon NIS-Elements imaging software were used to produce micrographs. Epifluorescence microscopy was used collect Z-stack image series from immunostained cultures shown in Fig. S4, S5 and Movie S3, S4. ImageJ and Matlab custom designed codes were used to create 3D surface plots (Fig. S4) and to reconstruct 3D neuronal morphologies (Fig. S5) based on the intensities of immunofluorescent signals.

### Network-wide activity mapping

Spontaneous extracellular AP were sampled across the entire microelectrode array by sequential scanning over 28 recording configurations. Up to 1024 randomly selected electrodes recorded neuronal activity in each configuration for 2 minutes. The average voltage traces, recorded by each electrode, were used to reconstruct the network-wide activity map (Fig. S6A). Since the largest extracellular APs occur near the AIS, and APs arising from axonal arbors have much smaller amplitude, regions with high-amplitude APs in the activity maps indicated the locations of the AISes^9,29, 30^.

### Electrical identification of individual neurons

Simultaneous access to signals arising from the AIS region enabled us to reconstruct the spatiotemporal distribution of extracellular APs, referred to as the ‘electrical footprint’ (Fig. S6C). To obtain these signals, we used high-density recording configurations covering the AIS locations revealed in the activity maps (Fig. S6A). In each configuration, blocks of 13×13 electrodes were connected to readout channels to sample neuronal activity for 2 minutes. Recorded signals were sorted by using the Spyking-Circus algorithm^55^, and electrical footprints were reconstructed by using custom-designed Matlab code. Because the first-occurring (initial) trace found in the electrical footprint colocalizes with the neuron’s AIS, it was used to trigger the averaging of voltage traces recorded across an entire axonal arbor (see below).

### Electrical imaging of axonal arbors

Array-wide averaging of voltage traces, synchronized with the initial trace (recorded from the AIS), reveals the spatiotemporal distribution of extracellular APs across an entire axonal arbor (Fig. S7 and Movie S5). The first step in obtaining these data was selecting 9 electrodes that were closest to the putative AIS. We next designed multiple recording configurations covering the entire array - in each configuration, 9 of the 1024 readout channels were routed to the 9 preselected electrodes, and remaining available channels were routed to randomly selected electrodes. Each configuration was used to sample neuronal activity during 2 minutes. Signals recorded by the 9 preselected electrodes in each configuration were sorted using the Spyking-Circus algorithm^55^. Timestamps of the sorted signals were used to trigger the averaging of voltage traces recorded across all other electrodes in the array. The spatiotemporal distribution of averaged signals was reconstructed using a custom-designed Matlab code.

### Electrical stimulation of individual neurons in the network

Stimulation performances of the HD-MEA system allow for stimulating any neuron in the culture with subcellular spatial precision^9,11, 30, 56^. Key concepts of the stimulation protocols that were used in this study are presented in Fig. S8. Extracellular electrical stimulations targeted at predefined axonal locations allow to experimentally control the direction of the axonal conduction^30^ (Fig. S8A). Electrical stimulation directed at the AIS was used to elicit orthodromic neuronal activation (Fig. S8A-left). Stimulation directed at distal axons was used to elicit antidromic neuronal activation (Fig. S8A-right). All stimulation protocols utilized balanced positive-first biphasic voltage pulses, with phase durations of 200 µs, because of their proven effectiveness in electrical stimulation^57^. Stimuli were applied to one electrode at a time at a frequency of 4 Hz. Neuronal responses to orthodromic stimulations were estimated by observing APs recorded from distal axons (Fig. S8A-left). Responses to antidromic stimulation were estimated by observing APs recorded from proximal axons and the AIS (Fig. S8A-right). To find optimal parameters for stimulation of individual neurons in the network, we used approaches thoroughly described in our previous study^30^. In brief, we applied neuron-wide stimulation over a range of voltages to reveal sites with the lowest activation threshold (Fig. S8B). Stimulation was applied at 4 Hz for voltages from ±10 to ±300 mV, with steps of ±10 mV. Each stimulation voltage was applied 60 times per site, and activation thresholds were defined as the minimum voltage to trigger an AP in 100% of the trials. To get more detailed excitability profiles, the most sensitive sites were then stimulated with voltages stepped by ±1 mV (Fig. S8C).

### Statistical analysis

All quantitative data presented in Fig. 7-9 are expressed as mean ± SEM. Numbers of biological replicates (N) used for analyses presented in Fig. 5-9 are denoted in the figures and are also stated in their legends. We used non-parametric tests for comparing distributions of parameters presented in Fig. 7-9, since normal distribution of the underlying data could not be determined unequivocally. The two-sided Mann-Whitney U-test was applied and a P-value < 0.05 was considered significant. Logarithmic scaling of graphs shown in Fig. 8B, C was used to display a wide range numerical data in a compact way.

## ACKNOWLEDGMENTS

We thank Olga Netsyk, Bryndis Birnir, Tanel Punga and Evgenii Bogatikov for help with experiments; Markus de Ruijter and Stefan Engblom for help with software developments; Marta K. Lewandowska and Yu-Fang Huang for critical discussions.

## ADDITIONAL INFOPRMATION

### Funding

This study was supported by Swedish Research Council grant 2016-02184, Swedish Research Council grant 2014-02048, Swedish Research Council grant 2014-07603, Swedish Research Council grant 2020-02040, Göran Gustafsson foundation for medical research, Hjärnfonden (the Swedish Brain Foundation), Olle Engkvist Byggmästare grant 197-0235, Margarethahemmets foundation and Bissen Brainwalk Foundation. The funders had no role in study design, data collection and analysis, decision to publish, or preparation of the manuscript.

### Author contributions

Conceptualization, experimental design, methodology, software development, visualization, validation, investigation, data analysis and writing were done by MR. The project supervision was coordinated by ARP.

### Data and materials availability

All data needed to evaluate the conclusions in the paper are present in the paper and the Supplementary Materials. There are no restrictions on materials used in the analysis. The datasets generated for this study, entitled ‘Electrical imaging of cortical and spinal axons using high-density microelectrode arrays’, are stored in Dryad (DOI: https://doi.org/10.5061/dryad.gxd2547r1).

## Supplementary Materials

### SUPPLEMENTARY FIGURES

**Figure S1 (related to Figure 2).**
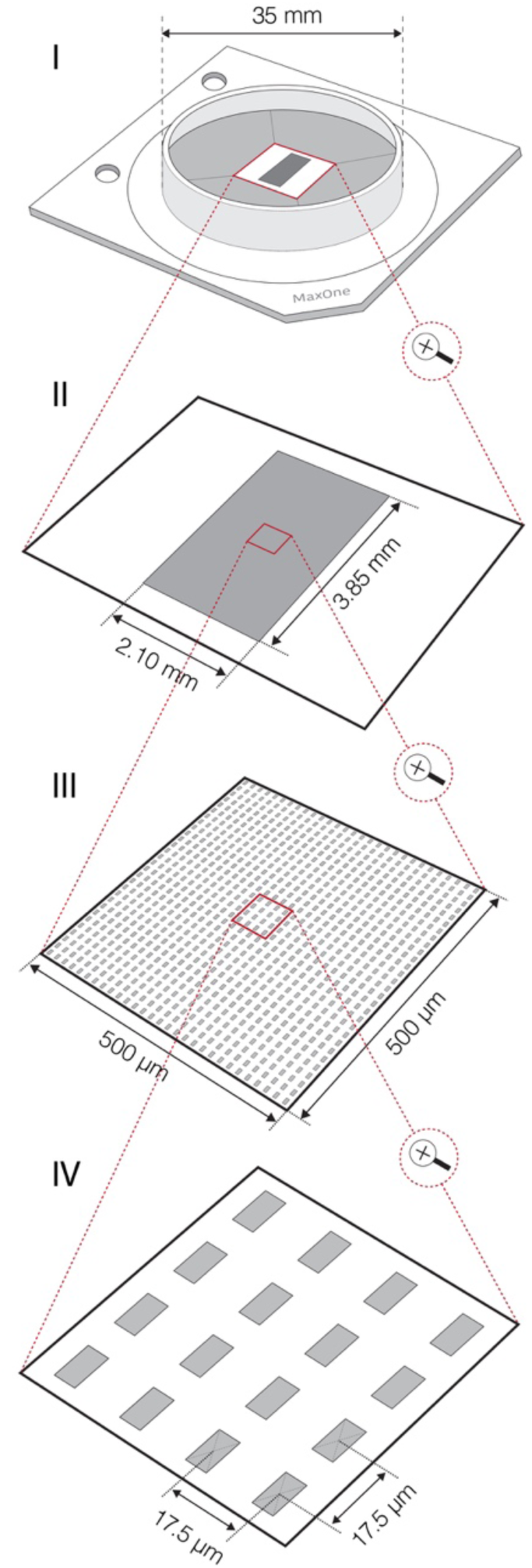
CMOS-based HD-MEA. Graphical scheme shows technical features of complementary-metal-oxide-semiconductor (CMOS)-based high-density (HD) microelectrode array (MEA) system that was used in this study. (I) Outlook of HD-MEA chip. (II) The MEA comprises 26,400 platinum electrodes packed within sensing a area of ∼8 mm^2^. Each of the electrodes can serve as a recording and/or stimulation source enabling bi-directional communication with any cell laying on the sensing area. (III) Dense arrangement of microelectrodes (3,264 electrodes per mm^2^) enables to access electrical activity of a single neuron across hundreds of sites at subcellular spatial resolution. (IV) Sizes of individual microelectrodes are 9.3×5.45 µm^2^; interelectrode center-to-center pitch is 17.5 µm. Up to 1024 readout channels can be simultaneously used to sample neuronal signals. Flexible switch matrix technology allows to route available channels to desired electrodes and to reconfigure the routing within few microseconds. Raw neuronal signals are recorded at 20-kHz sampling rate.

**Figure S2 (related to Figure 2).**
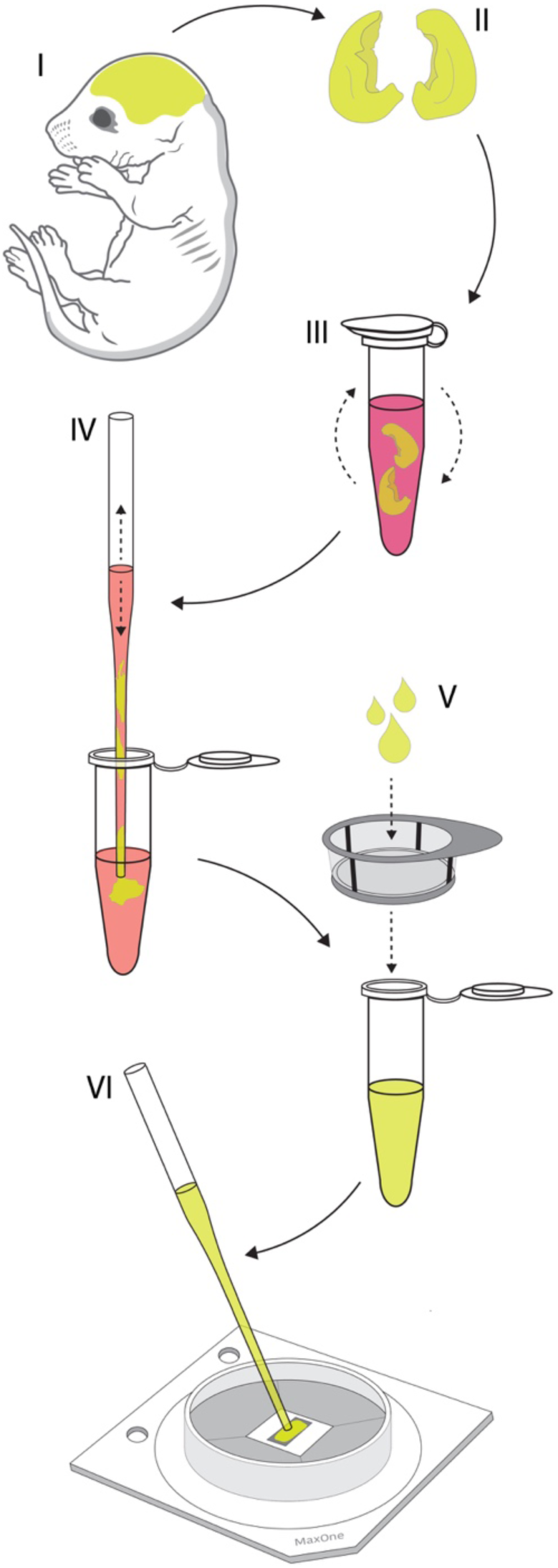
Culturing primary cortical neurons on the HD-MEA. Graphical illustration shows the key steps of the protocol used for culturing rat’s primary cortical neurons on the HD-MEA surface. (I) Entire brains were obtained from Sprague Dawley rat embryos dissected on embryonic day 18. (II) Cortices were isolated from the brains and cortical meninges were removed. (III) Cortices were dissociated enzymatically in trypsin supplemented with 0.25% EDTA. (IV) Cortices were dissociated physically by trituration in the plating medium. (V) Suspension of dissociated cells was filtered using 40-µm cell strainer to remove cell aggregates and debris. (VI) Filtered cells were plated directly on the chip’s sensing area coated by a thin layer of laminin. Cells were seeded in a 20 µl drop containing ∼50,000 cells in total.

**Figure S3 (related to Figure 2).**
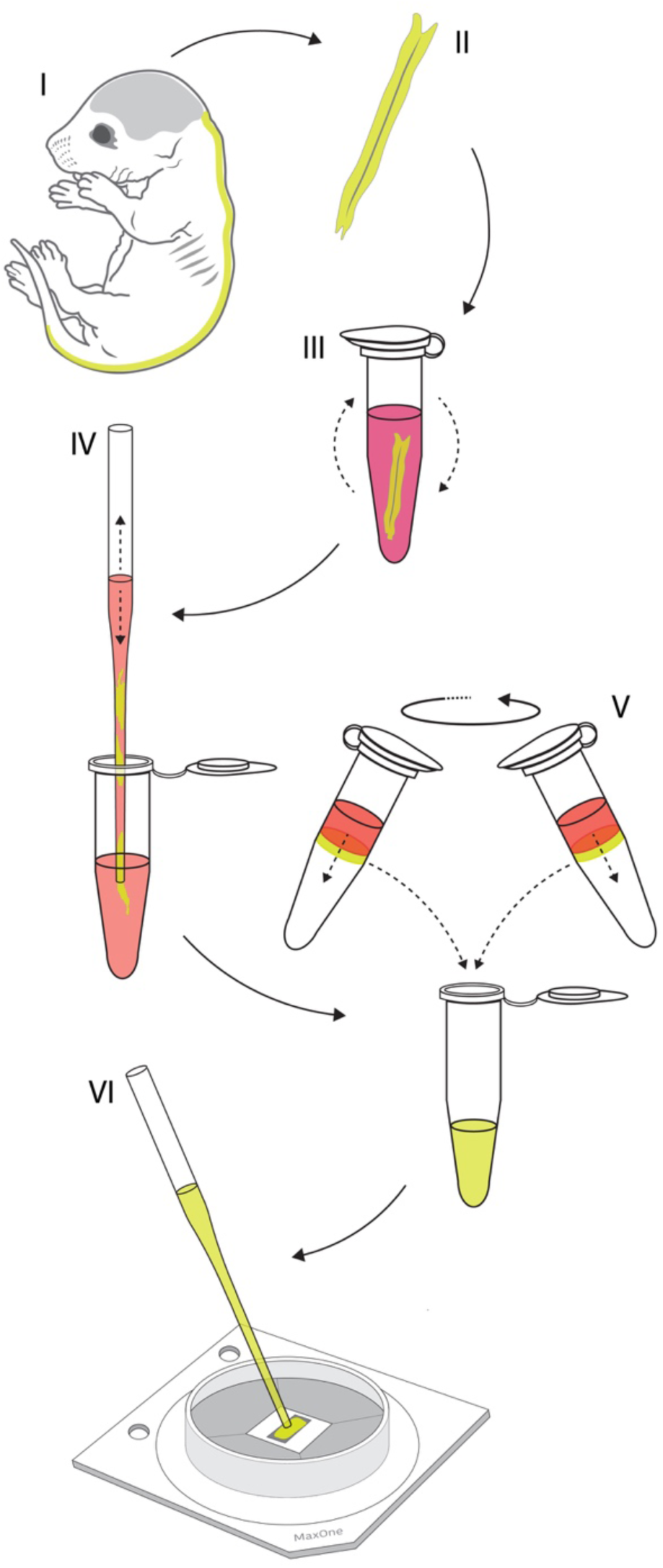
Culturing primary motor neurons on the HD-MEA. Graphical illustration shows key steps of the protocol used for culturing rat’s primary motor neurons on HD-MEA surface. (I) Entire spinal cords were isolated from Sprague Dawley rat embryos dissected on embryonic day 14. (II) Dorsal root ganglia were detached and meninges were removed from isolated spinal cords. (III) Spinal cords were dissociated enzymatically in trypsin supplemented with 0.25% EDTA. (IV) Spinal cords were next dissociated physically by trituration in the plating medium. Suspension of dissociated cells was filtered using 40 µm cell strainer to remove cell aggregates and debris. (V) Density gradient centrifugation was used to fractionate motor neurons from the filtered cell-suspension. (VI) Isolated cells were plated directly on the chip’s sensing area coated by a thin layer of laminin. Cells were seeded in a 20 µl drop containing ∼50,000 cells in total.

**Figure S4 (related to Figure 2).**
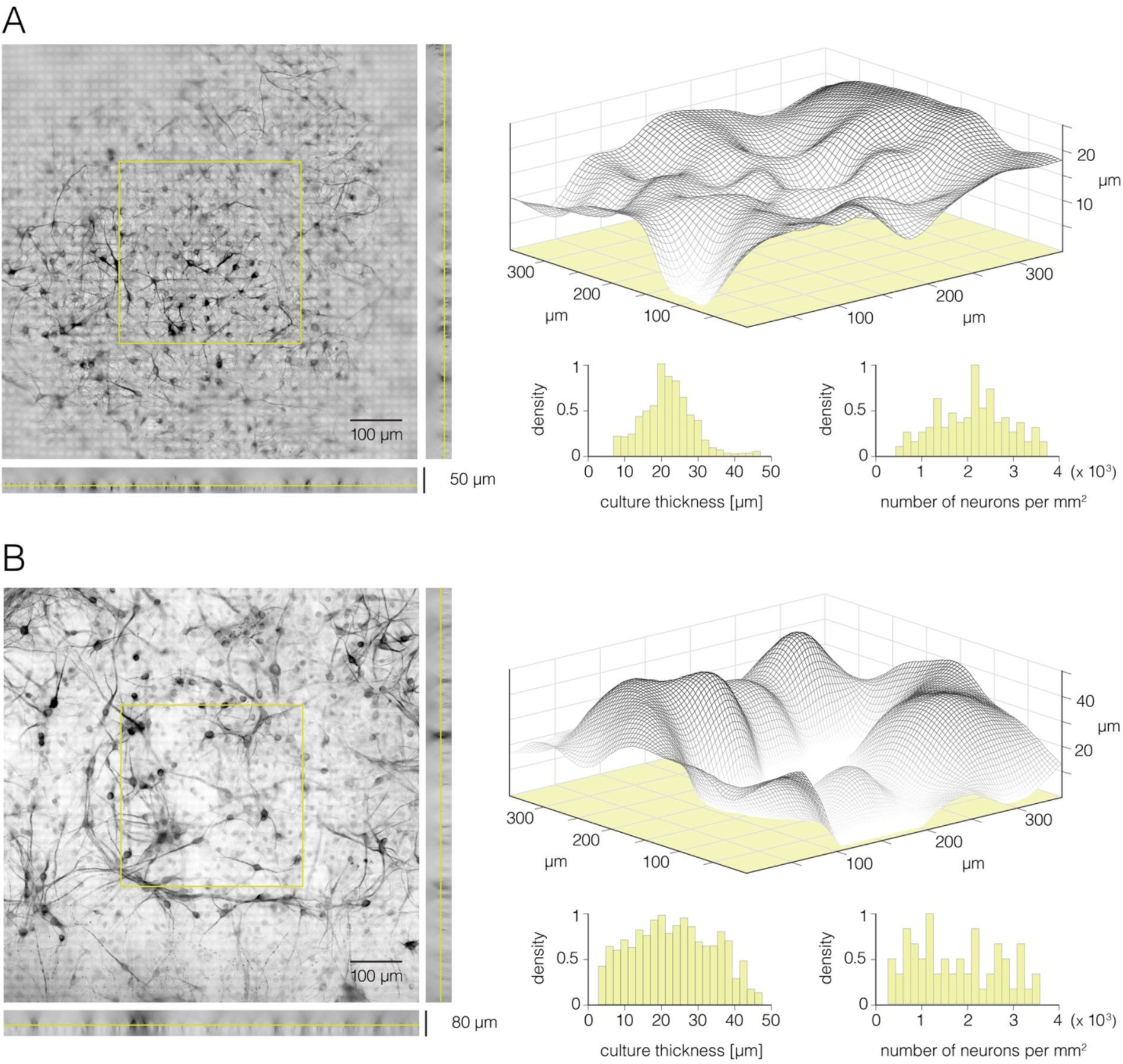
Thickness and cell-density of neuronal cultures grown on the HD-MEA. **(A)** Rat primary cortical culture grown on the HD-MEA surface – 16 DIV. (Left) Micrograph of the cortical culture immunostained against β-III tubulin. Epifluorescence microscopy was used to collect Z-stack image series. Z-position of the focus plane shown is marked on the side view projections (yellow lines). Yellow square superimposed over the center of the micrograph denotes a section of the Z-stack data that was used to produce the graph shown on the right. (Right) Surface plot reconstructed from the Z-stack data denoted in the left panel. Three-dimensional graph (gray net) displays intensities of fluorescent signals. Reconstructed graph was used to estimate thickness of the culture. Distribution of local thicknesses and cell-densities sampled over discrete areas of 100×100 µm^2^ are presented in the histograms bellow. **(B)** Rat’s primary motor-neuron culture grown on HD-MEA surface – 12 DIV. (Left) Micrograph of the motor-neuron culture immunostained against β-III tubulin. Epifluorescence microscopy was used to collect Z-stack image series. Z-position of the focus plane shown is marked on the side view projections (yellow lines). Yellow square superimposed over the center of the micrograph denotes a section of the Z-stack data that was used to produce the graph shown on the right. (Right) Surface plot reconstructed from the Z-stack data denoted in the left panel. Three-dimensional graph (gray net) displays intensities of fluorescent signals. Reconstructed graph was used to estimate thickness of the culture. Distribution of local thicknesses and cell-densities sampled over discrete areas of 100×100 µm^2^ are presented in two histograms below.

**Figure S5 (related to Figure 2).**
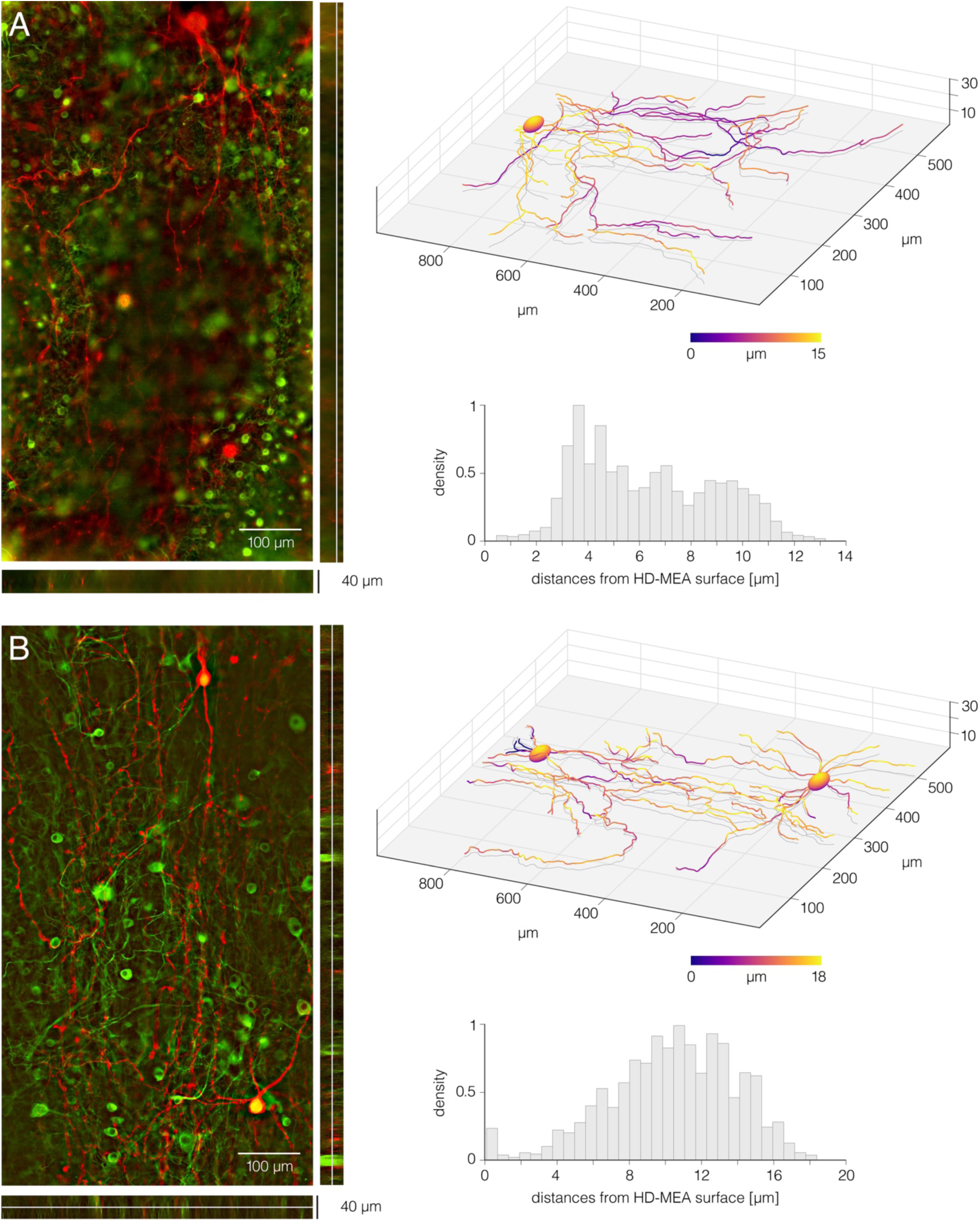
Disposition of cortical and spinal neurites with respect to the HD-MEA surface. **(A)** Cortical neurons grown on the HD-MEA surface – 16 DIV. (Left) Micrograph of fluorescently labelled cortical neurons. Neurons were sparsely transfected at 14 DIV using RFP-expressing plasmid (pAAV-hSyn-mCherry); the RFP-expression was optically confirmed in 6 neurons. The culture was fixed at 16 DIV and immunostained against beta-III tubulin to label all neurons in the culture (green); the culture was also immunostained against RFP to intensify fluorescent signal in the transfected neurons (red). Epifluorescence microscopy was used to collect Z-stack image series. (Right-up) Three-dimensional neuronal morphology reconstructed from optical data shown in the left panel. Local distances of neuronal processes from the HD-MEA surface are color-coded. (Right-down) Histogram shows distribution of local distances of neuronal processes from the HD-MEA surface. **(B)** Motor neurons grown on the HD-MEA surface – 12 DIV. (Left) Micrograph of fluorescently labelled motor neurons. Individual neurons were sparsely transfected at 10 DIV using RFP-expressing plasmid (pAAV-hSyn-mCherry); the RFP-expression was optically confirmed in 14 neurons. The culture was fixed at 12 DIV and immunostained against beta-III tubulin to label all neurons in the culture (green); the culture was also immunostained against RFP to intensify fluorescent signal in the transfected neurons (red). Epifluorescence microscopy was used to collect Z-stack image series. (Right-up) Three-dimensional neuronal morphologies reconstructed from optical data shown in the left panel. Local distances of neuronal processes from the HD-MEA surface are color-coded. (Right-down) Histogram shows distribution of local distances of neuronal processes from the HD-MEA surface. ImageJ and Matlab custom designed codes were used to reconstruct three-dimensional neuronal morphologies shown here.

**Figure S6 (related to Figure 2).**
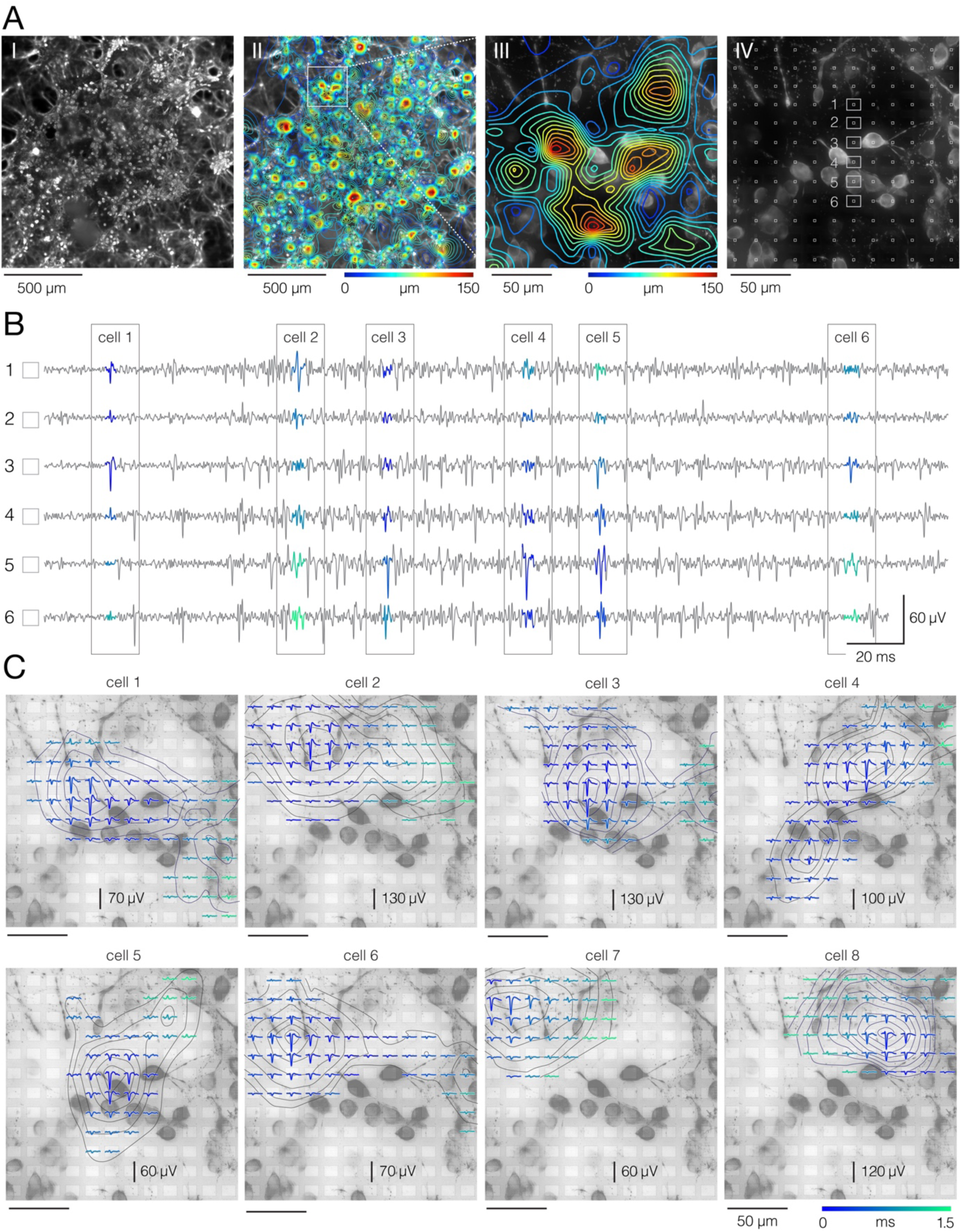
Electrical identification of individual neurons in the culture. **(A)** Network-wide electrical activity. (I) Micrograph of fluorescently labeled cortical culture grown on the HD-MEA. The culture was fixed at 16 DIV and immunostained against Map2. Epifluorescence microscopy was used to collect Z-stack image series (presented in Movie S3). Herein, presented is maximum signal intensity. (II) Network-wide activity map. Spontaneous APs were sampled across all microelectrodes for 2 minutes. Average amplitudes of recorded signals were used to reconstruct the map; signal amplitude is color-coded. Micrograph of the corresponding culture is set in back. White square superimposed over the map frames the region magnified in III. (IV) Positions of the recording microelectrodes (white squares in front) with respect to locations of neuronal somas (micrograph in the background); same region as in III; Z-stack image series are presented in Movie S4. Voltage traces recorded from six denoted electrodes (1-6) are shown in (B). **(B)** Discerning between APs recorded from neighboring neurons. Spike-sorting algorithms enable discerning mixed signals recorded from neighboring neurons and thereby extracting APs arising from individual neurons. Voltage traces recorded from 6 electrodes (denoted in A-IV) are presented in gray. Sorted signals from different cells (1-6) are colored. **(C)** Reconstructed electrical activities of individual neurons. Averaging of sorted signals over adjacent electrodes reveals the spatiotemporal distribution of a single neuron’s activity. Such signal representation is called an “extracellular AP footprint”. Herein, presented are electrical footprints of 8 neighboring neurons, superimposed over micrograph of the corresponding culture (same as in A-IV). Temporal distribution of APs detected on different electrodes is color-coded.

**Figure S7 (related to Figure 2).**
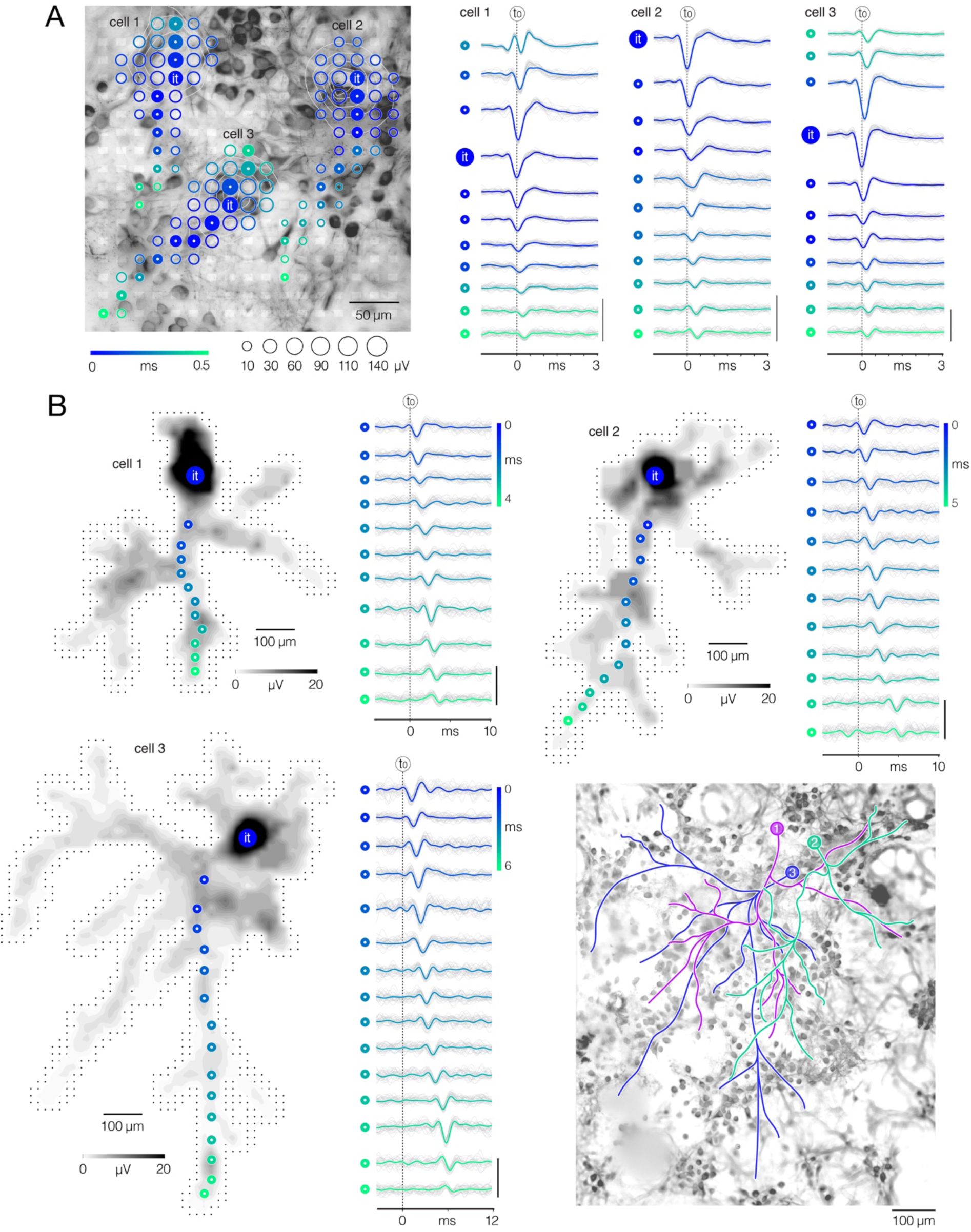
Electrical imaging of axonal arbors. **(A)** Electrical footprints reveal proximal neuronal compartments. (Left) Electrical footprints of three neighboring cortical neurons (denoted as cell 1-3). Circle sizes indicate average amplitudes of APs detected on individual microelectrodes; occurrence time of detected AP peaks relative to timing of the initial trace (t_0_) is color-coded. Color-filled circles denote locations along putative dendrite-AIS-axon axis; APs recorded from the denoted locations are shown in right. Micrograph of the corresponding culture is shown in the back; the culture was fixed at 16 DIV and immunostained against Map2. White semitransparent rectangles superimposed over the micrograph represent microelectrodes. (Right) AP waveforms recorded along dendrite-AIS-axon axis as denoted in left. Action-potential waveforms are temporally aligned with respect to occurrence time of the initial trace (it); note temporal shift of the aligned AP peaks relative to t_0_ (black-dashed line in the back). Average AP waveforms are colored (blue-cyan) and superimposed over the corresponding single-trials (gray). Color-coding same as in left. Signal scalebar: 100 µV. **(B)** Spike-trigger averaging of sorted APs reveals entire axonal arbors. (Left-up, Right-up, Left-down) Spatial distribution of axonal APs recorded across an entire HD-MEA; same neurons as in (a); AP amplitude is color-coded (gray scale). It - initial trace; color-filled circles denote recording sites for axonal APs shown on the side. Black dots in the background represent microelectrodes. Axonal AP waveforms recorded from the denoted sites are presented on the side; their occurrence time is color-coded. Average AP waveforms are colored (blue-cyan) and superimposed over the corresponding single-trials (gray). Signal scalebar: 100 µV. (Right-down) Axonal contours of the three cortical neurons (denoted as 1-3) superimposed over the micrograph of the corresponding culture. Spike-trigger averaged signals of the three neurons is presented in Movie S5. Axonal contours are estimated by observing spatial movements of the signal peaks in consecutive movie frames. The culture was fixed at 16 DIV and immunostained against Map2. Epifluorescence microscopy was used to collect Z-stack image series (presented in Movie S3).

**Figure S8 (related to Figure 6).**
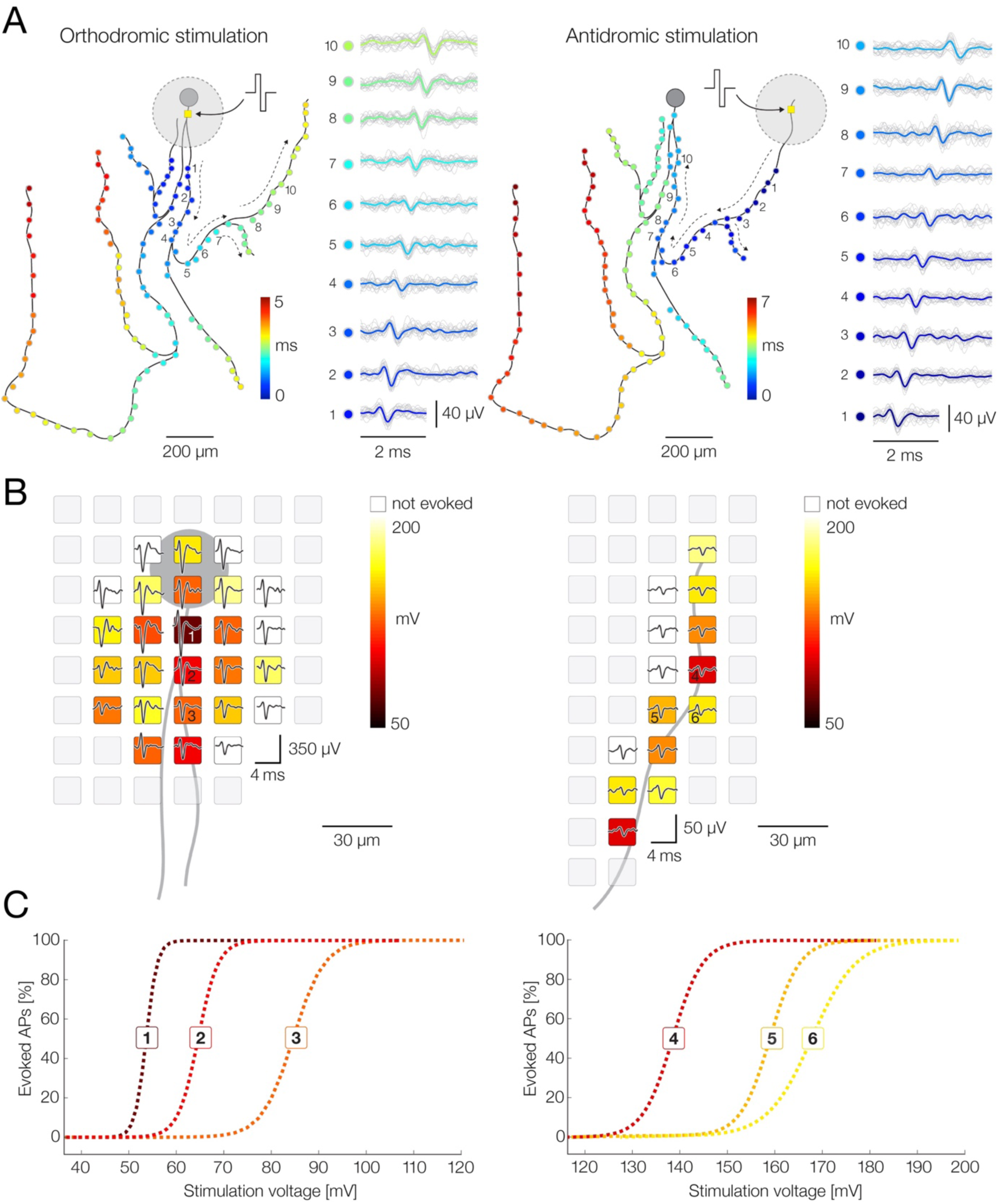
Targeted microstimulation of axons using the HD-MEA system. **(A)** Schematic representation of electrical stimulations directed at proximal (left) and distal (right) axon of a motor neuron. Stimulation-triggered axonal APs are presented through colored circles superimposed over black axonal contour; conduction time of the presented APs is color-coded. Black dashed arrows indicate directions of axonal conduction. Waveforms of axonal APs recorded from ten denoted locations (1-10) are shown on the sides; averaged waveforms (color-coded traces) are superimposed over single recording trials (pale-gray traces). Positive-first biphasic voltage pulses were used to stimulate the neuron. Stimulation sites are denoted by yellow squares centered within large gray circles; the gray circles represent artifacts induced by the stimulations. (Left) Electrical stimulation targeted at proximal axon initiated an orthodromic signal conduction; stimulation voltage amplitude of ±55 mV was used to stimulate the neuron 60 times at 4 Hz. (Right) Electrical stimulation targeted at distal axon initiated an antidromic signal conduction; stimulation voltage amplitude of ±150 mV was used to stimulate the neuron 60 times at 4 Hz. **(B)** Identification of stimulation electrodes and voltage magnitudes for orthodromic (left) and antidromic (right) stimulation. Average waveforms of neuronal APs (black traces) recorded from proximal and distal axons are superimposed over corresponding “stimulation maps” – spatial distribution of identified stimulation thresholds for effective activation of the neuron. Stimulation thresholds are color-coded and are assigned to physical location of stimulation electrodes. (Left) Stimulation map obtained near proximal axon. Note: the lowest stimulation threshold colocalized with the neuronal signal with the largest amplitude (putative AIS). (Right) Stimulation map obtained near the distal axon. Note larger values of stimulation thresholds as compared with the proximal stimulation map. (left-right) Stimulation electrodes with the lowest thresholds were used for orthodromic and antidromic stimulations presented in a. Excitability profiles for six denoted electrodes (1-6) are shown in c. **(C)** Excitability profiles of six electrodes with the lowest thresholds for orthodromic (1-3) and antidromic (4-6) stimulation. Each of the electrodes was probed with range of stimulation voltages (±10 to ±300 mV) and stimulation-triggered neuronal response were used to express the excitability profiles. Each of the voltages was applied 100 times per electrode. Stimulation thresholds (color-coded in b) were defined as the minimal voltages that activated the neuron in 100 % of the stimulation trials.

### SUPPLEMENTARY MOVIES

**Movie S1 (related to Figure 1). Functional morphologies of cortical axons displayed in real-time. (Left)** Reconstructed functional morphologies of cortical axons superimposed over a micrograph of the corresponding culture. Morphologies of individual neurons are presented through consecutive sequences of AP propagation paths as traced across axonal arbors in real-time. Functional morphologies of 15 neurons are displayed successively and stacked on top of each other; neuron count is color-coded; total length of reconstructed axons amounts 348.82 mm. A micrograph of immunostained rat’s primary cortical culture grown directly on the HD-MEA surface is shown in the background. The culture was fixed immediately after the acquisition of electrophysiological data, and fluorescently labelled against β-III tubulin at 16 DIV. Epifluorescence microscopy was used to collect optical data. **(Right)** Spontaneous electrical activities of cortical neurons as monitored across an entire HD-MEA. Amplitudes of electrical signals recorded by individual electrodes are color-coded in grey scale; recorded signals are mapped spatially across all microelectrodes and temporally over discrete timeframes to produce the movie. Trajectories of axonal signals are traced over consecutive timeframes to reconstruct morphologies of individual neurons. Time-gap between consecutive timeframes is 50 µs. Real conduction times of traced signals are color-coded and presented through reconstructed morphologies (blue-cyan scale). Functional morphologies of 15 neurons are traced successively and simultaneously shown on both right and left panel.

**Movie S2 (related to Figure 1). Functional morphologies of spinal axons displayed in real-time. (Left)** Reconstructed functional morphologies of motor neuron axons superimposed over a micrograph of the corresponding culture. Morphologies of individual neurons are presented through consecutive sequences of AP propagation paths as traced across axonal arbors in real-time. Functional morphologies of 15 neurons are displayed successively and stacked atop each other; neuron count is color-coded; total length of reconstructed axons amounts 282.72 mm. Micrograph of immunostained rat’s primary motor neuron culture grown directly on the HD-MEA surface is shown in the background. The culture was fixed immediately after the acquisition of electrophysiological data, and fluorescently labelled against β-III tubulin at 12 DIV. Epifluorescence microscopy was used to collect optical data. **(Right)** Spontaneous electrical activities of motor neurons as monitored across an entire HD-MEA. Amplitudes of electrical signals recorded by individual electrodes are color-coded in grey scale; recorded signals are mapped spatially across all microelectrodes and temporally over discrete timeframes to produce the movie. Trajectories of axonal signals are traced over consecutive timeframes to reconstruct morphologies of individual neurons. Time-gap between consecutive timeframes is 50 µs. Real conduction times of traced signals are color-coded and presented through reconstructed morphologies (blue-cyan scale). Functional morphologies of 15 neurons are traced successively and simultaneously shown on both right and left panel.

**Movie S3 (related to Figure 2). Cortical culture grown on HD-MEA surface – 20x magnification.** Z-stack image series of cortical culture grown on the HD-MEA. The presented culture-section covered an area of 1.3×1.3 mm^2^ and interfaced with 5,476 electrodes. The culture was fixed at 16 DIV and immunostained against Map2. Epifluorescence microscopy was used to collect Z-stack image series. Data obtained from the same culture is presented in Fig. S6, S7 and in Movie S4.

**Movie S4 (related to Figure 2). Cortical culture grown on HD-MEA surface – 40x magnification.** Z-stack image series of cortical culture grown on the HD-MEA. The presented culture-section covered an area of 0.3×0.3 mm^2^ and interfaced with 289 electrodes. The culture was fixed at 16 DIV and immunostained against Map2. Epifluorescence microscopy was used to collect Z-stack image series. Data obtained from the same culture is presented in Fig. S6, S7 and in Movie S3.

**Movie S5 (related to Figure 2). Electrical imaging of axonal arbors.** Spatiotemporal distribution of extracellular APs as propagating across axons of three cortical neurons. Amplitudes of APs recorded by individual electrodes are color-coded; recorded signals are mapped spatially across all microelectrodes and temporally over discrete timeframes to produce the movie. Time-gap between consecutive timeframes is 50 µs. Data obtained from same neurons is presented in Fig. S7. Micrograph of the corresponding culture is shown in the back; the culture was fixed at 16 DIV and immunostained against Map2. Epifluorescence microscopy was used to collect optical data. Z-stack image series of the same culture-section is presented in Movie S3.

**Movie S6 (related to Figure 3). Mapping signals across axonal arbor – step 1.** Simple planar threshold set to 9 STD of the estimated noise enables detection of high-amplitude AP peaks along the axonal arbor. **(Left)** 3D graph shows planar threshold (red semitransparent plane) applied on a section of neuronal signal (3D gray-lined mesh) reconstructed over consecutive timeframes. 2D profiles of the reconstructed signal are projected on the graph’s side planes (gray-filled hills). Signal cutouts found above the threshold are projected on the graph’s bottom (red-line bordered patches). Detected signal peaks are denoted by green circles projected perpendicularly from top to bottom of the graph. Axonal contour shown at the graph’s bottom was estimated by observing spatial movements of the signal peaks over consecutive timeframes. **(Right)** 2D representation of the data shown in left. Spatiotemporal distribution of the detected peaks (blue-cyan circles) superimposed over axonal contour (same as in left). Peaks detected over consecutive timeframes are denoted by green circles; detected peaks are embedded within 2D projection of the signal cutouts found above the threshold (red-line bordered patches, same as in left). Occurrence time of detected peaks is color-coded (blue-cyan). Neuronal electrical activity reconstructed over consecutive timeframes is shown in the background; average AP amplitude is color-coded (gray scale). Section of the same data is presented in Fig. 3A.

**Movie S7 (related to Figure 3). Mapping signals across axonal arbor – step 2.** Adaptations of the threshold based on spatial and temporal coordinates of previously mapped peaks enables detection of low-amplitude APs along the axonal arbor. **(Left)** 3D graph shows local thresholds (red semitransparent circles) applied on a section of neuronal signal (3D gray-lined mesh) reconstructed over consecutive timeframes. Local thresholds are centered on XY coordinates of previously detected peaks and are adapted to detect neighboring peaks in preceding and succeeding timeframes; detection fields of local thresholds are 50 µm in radius, and are set to 2 STD of the estimated noise. Signal cutouts found above the threshold are projected on the graph’s bottom (red-line bordered patches). Detected signal peaks are denoted by green circles projected perpendicularly from top to the bottom of the graph. Axonal contour shown at the graph’s bottom was estimated by observing spatial movements of the signal peaks over consecutive timeframes. **(Right)** 2D representation of the data shown in left. Spatiotemporal distribution of the detected peaks superimposed over axonal contour. Peaks detected over consecutive timeframes are denoted by green circles appearing in each frame; detected peaks are embedded within 2D projection of the signal cutouts found above the threshold (red-line bordered patches, same as in left). Previously detected peaks are denoted by white-filled circles; newly detected peaks are denoted by accumulating blue-cyan circles. Occurrence time of detected peaks is color-coded (blue-cyan). Neuronal electrical activity reconstructed over consecutive timeframes is shown in the background; average AP amplitude is color-coded (gray scale). Section of the same data is presented in Fig. 3B.

**Movie S8 (related to Figure 3). Mapping signals across axonal arbor – step 3.** Adaptations of the threshold based on spatial and temporal coordinates of previously mapped peaks enables detection of low-amplitude APs along axonal arbor. (Left) 3D graph shows local thresholds (red semitransparent circles) applied on a section of neuronal signal (3D gray-lined mesh) reconstructed over consecutive timeframes. Local thresholds are centered on XY coordinates of previously detected peaks and are adapted to detect neighboring peaks in preceding and succeeding timeframes; detection fields of local thresholds are 100 µm in radius, and are set to 1 STD of the estimated noise. Signal cutouts found above the threshold are projected on the graph’s bottom (red-line bordered patches). Detected signal peaks are denoted by green circles projected perpendicularly from top to the bottom of the graph. Axonal contour shown at the graph’s bottom was estimated by observing spatial movements of the signal peaks over consecutive timeframes. (Right) 2D representation of the data shown in left. Spatiotemporal distribution of the detected peaks superimposed over axonal contour. Peaks detected over consecutive timeframes are denoted by green circles appearing in each frame; detected peaks are embedded within 2D projection of the signal cutouts found above the threshold (red-line bordered patches, same as in left). Previously detected peaks are denoted by white-filled circles; newly detected peaks are denoted by accumulating blue-cyan circles. Occurrence time of detected peaks is color-coded (blue-cyan). Neuronal electrical activity reconstructed over consecutive timeframes is shown in the background; average AP amplitude is color-coded (gray scale). Section of the same data is presented in Fig. 3C.

**Movie S9 (related to Figure 4). Direct interconnection of mapped signal peaks.** Fragments of axonal conduction trajectory reconstructed based on spatiotemporal proximities of axonal signal peaks. **(Left)** Closest signal peaks found within 100 µm Euclidean range (moving white circles) and mapped over consecutive timeframes are interconnected via direct links (transient white lines). Signal peaks found in a current timeframe are denoted by white-filled circles. Signal peaks found in three preceding timeframes are denoted by colored circles (blue-cyan); colors encode signal conduction time. Direct links established in preceding timeframes are presented by thin black lines; direct links outline fragments of axonal conduction trajectory. Non-connected peaks are denoted by black-filled circles. Spontaneous activity of the corresponding neuron is shown in the background; average amplitude of the axonal signal is color-coded (gray scale). **(Right)** Fragments of axonal conduction trajectory revealed by direct interconnection (as shown in Left) are presented by thick colored lines (blue-cyan); colors encode conduction time of axonal signals. Spontaneous activity of the corresponding neuron is shown in the background; average amplitude of the axonal signal is color-coded (gray scale). Section of the same data is presented in Fig. 4A.

**Movie S10 (related to Figure 4). Skeletonization-assisted interconnection of mapped signal peaks.** Skeletonized axonal signals mediate interconnection of mapped peaks that could not be interconnected directly. **(Left)** Fragments of axonal conduction trajectory revealed by direct interconnection (as shown in Movie S9) are presented through dark-gray contour. Remaining gaps in the trajectory are filled via skeletonization-assisted interconnection of mapped peaks. Signal peaks found within 200-µm Euclidean range (moving white circles) and mapped in two consecutive timeframes are interconnected via remnants of the skeletonized signal (irregular white lines) - signal averaged over the two consecutive timeframes (Δt = 100 µs) is skeletonized to infer directionality of the axonal conduction. Signal peaks found in a current timeframe are denoted by white-filled circles. Signal peaks found in the preceding timeframe are denoted by colored circles (blue-cyan); colors encode timing of the signal conduction. Non-connected peaks are denoted by black-filled circles. Spontaneous activity of the corresponding neuron is shown in the background; average amplitude of the axonal signal is color-coded (gray scale). **(Right)** Axonal conduction trajectory revealed by direct and skeletonization-assisted interconnection (as shown in Left) are presented by thick colored lines (blue-cyan); colors encode timing of the signal conduction. Spontaneous activity of the corresponding neuron is shown in the background; average amplitude of the axonal signal is color-coded (gray scale). Section of the same data is presented in Fig. 4B.

**Movie S11 (related to Figure 4). Indirect interconnection of mapped signal peaks.** Indirect interconnection enables to interlink spatiotemporally distanced peaks that could not be interconnected via direct nor skeletonization-assisted interconnection. **(Left)** Axonal conduction trajectory revealed by direct and skeletonization-assisted interconnection (as shown in Movie S10) are presented through dark-gray contour. Remaining gaps in the trajectory are filled via indirect interconnection of mapped peaks. Spatiotemporally distanced peaks that are mapped in every second timeframe and found within 400 µm Euclidean range (moving white circles) are interconnected indirectly - signal averaged over three consecutive timeframes (Δt = 150 µs) is skeletonized to interconnect peaks mapped in 1^st^ (current) and 3^rd^ (second preceding) timeframe (irregular white lines); data for 2^nd^ (first preceding) timeframe is predicted based on the average conduction velocity observed in previously reconstructed fragments of the trajectory. Signal peaks found in a current timeframe are denoted by white-filled circles. Signal peaks found in the second preceding timeframe are denoted by colored circles (blue-cyan); colors encode timing of the signal conduction. Data predicted for the first preceding timeframe is denoted by framed white-filled circles. Spontaneous activity of the corresponding neuron is shown in the background; average amplitude of the axonal signal is color-coded (gray scale). **(Right)** Axonal conduction trajectory revealed by direct, skeletonization-assisted and indirect interconnection (as shown in Left) are presented by thick colored lines (blue-cyan); colors encode timing of the signal conduction. Spontaneous activity of the corresponding neuron is shown in the background; average amplitude of the axonal signal is color-coded (gray scale). Section of the same data is presented in Fig. 4C.

**Movie S12 (related to Figure 6). Stimulation-aided reconstruction of axonal conduction trajectories.** Axonal conduction trajectories obtained from stimulation-triggered activities of a single motor neuron. Electrical stimulation was directed at three predefined axonal sites (yellow semi-transparent circles) – the AIS (stimulation I), distal axon on the right (stimulation II) and distal axon on the left (stimulation III). We used charge-balanced positive-first biphasic voltage pulses with phase durations of 200 µs and amplitudes of ±30, ±50 and ±40 mV for the stimulation I, II and III, respectively. Each of the axonal sites was stimulated 60 times at 4 Hz. Stimulation sites and amplitudes used specifically for this experiment were previously optimized (see Fig. S8). Stimulation-triggered activities were reconstructed and averaged across an entire array. Reconstructed activities were mapped spatially across all microelectrodes and temporally over discrete timeframes to produce the movie. Time-gap between consecutive timeframes is 50 µs. Average signal amplitude is color-coded (gray scale). Observing the spatial movements of signal peaks in consecutive movie frames enabled us to visually track axonal AP conduction in different directions and to reconstruct axonal conduction trajectories. Conduction times of stimulation-triggered APs are color-coded (blue-cyan scale). Conduction trajectories of the same neuron are presented in Fig. 6A and in Movie S13.

**Movie S13 (related to Figure 6). Visually tracked axonal conduction trajectories.** Axonal conduction trajectories obtained from spontaneous and stimulation-triggered activities of a single motor neuron (same neuron as in Movie S12). Spontaneous neuronal activity was mapped spatially across all microelectrodes and temporally over discrete timeframes to produce the movie shown in the background. Time-gap between consecutive timeframes is 50 µs. Average signal amplitude is color-coded (gray scale). Observing the spatial movements of signal peaks in consecutive movie frames enabled us to visually track axonal AP conduction and to reconstruct axonal conduction trajectories. Conduction times of tracked APs are color-coded (blue-cyan scale). Conduction trajectories of the same neuron are presented in Fig. 6A and in Movie S12.

**Movie S14 (related to Figure 7). Functional morphologies of cortical and spinal axons.** Functional morphologies of cortical (left) and spinal (right) axonal arbors reconstructed based on spontaneous neuronal activities. Branching orders of reconstructed neurites are color-coded (brown scale); neuronal somas are presented by black-filled circles. Spontaneous activities of corresponding neurons are displayed in the background. Amplitudes of electrical signals recorded by individual electrodes are color-coded in grey scale; recorded signals are mapped spatially across all microelectrodes and temporally over discrete timeframes to produce the movie. Propagation time of axonal APs is displayed in upper-left corner. The time gap between consecutive timeframes is 50 µs. Functional morphologies of the two neurons are also presented in Fig. 7A.

**Movie S15 (related to Figure 8). Action potential waveforms as traced across axonal functional morphologies.** Examples of AP waveforms extracted from functional morphologies of cortical (left) and motor-neuron (right) axons. **(Left)** Functional morphology of a cortical axon (white axonal contour) is displayed over spontaneous activity of the corresponding neuron; neuronal soma is presented by black-filled circle. AP waveforms traced across selected axonal path are displayed beside; average AP amplitudes are color-coded. **(Right)** Functional morphology of a motor neuron axon (white axonal contour) is displayed over spontaneous activity of the corresponding neuron; neuronal soma is presented by black-filled circle. AP waveforms traced across selected axonal path are displayed beside; average AP amplitudes are color-coded. **(Middle)** Note difference in scalebars for AP waveforms obtained from proximal and distal axons. Functional morphologies of the two neurons are also presented in Fig. 8A.

## REFERENCES

1 Stifani, N. Motor neurons and the generation of spinal motor neuron diversity. Frontiers in cellular neuroscience 8, 293 (2014).

2 Hodgkin, A. L. & Huxley, A. F. A quantitative description of membrane current and its application to conduction and excitation in nerve. The Journal of physiology 117, 500 (1952).

3 Alcami, P. & El Hady, A. Axonal computations. Frontiers in Cellular Neuroscience 13, 413 (2019).

4 Shu, Y., Hasenstaub, A., Duque, A., Yu, Y. & McCormick, D. A. Modulation of intracortical synaptic potentials by presynaptic somatic membrane potential. Nature 441, 761–765 (2006).

5 Alle, H. & Geiger, J. R. Analog signalling in mammalian cortical axons. Current opinion in neurobiology 18, 314–320 (2008).

6 Alle, H. & Geiger, J. R. Combined analog and action potential coding in hippocampal mossy fibers. Science 311, 1290–1293 (2006).

7 Zbili, M. & Debanne, D. Past and future of analog-digital modulation of synaptic transmission. Frontiers in cellular neuroscience 13, 160 (2019).

8 Geiger, J. R. & Jonas, P. Dynamic control of presynaptic Ca2+ inflow by fast-inactivating K+ channels in hippocampal mossy fiber boutons. Neuron 28, 927–939 (2000).

9 Radivojevic, M. et al. Tracking individual action potentials throughout mammalian axonal arbors. Elife 6, e30198 (2017).

10 Debanne, D. Information processing in the axon. Nature Reviews Neuroscience 5, 304–316 (2004).

11 Bakkum, D. J. et al. Tracking axonal action potential propagation on a high-density microelectrode array across hundreds of sites. Nature communications 4, 1–12 (2013).

12 Debanne, D., Campanac, E., Bialowas, A., Carlier, E. & Alcaraz, G. Axon physiology. Physiological reviews 91, 555–602 (2011).

13 Saliani, A. et al. Axon and myelin morphology in animal and human spinal cord. Frontiers in neuroanatomy 11, 129 (2017).

14 Faisal, A. A. & Laughlin, S. B. Stochastic simulations on the reliability of action potential propagation in thin axons. PLoS computational biology 3, e79 (2007).

15 Faisal, A. A., White, J. A. & Laughlin, S. B. Ion-channel noise places limits on the miniaturization of the brain’s wiring. Current Biology 15, 1143–1149 (2005).

16 White, J. A., Rubinstein, J. T. & Kay, A. R. Channel noise in neurons. Trends in neurosciences 23, 131–137 (2000).

17 Neishabouri, A. & Faisal, A. A. Axonal noise as a source of synaptic variability. PLoS computational biology 10, e1003615 (2014).

18 Skaugen, E. & Walløe, L. Firing behaviour in a stochastic nerve membrane model based upon the Hodgkin—Huxley equations. Acta Physiologica Scandinavica 107, 343–363 (1979).

19 Schneidman, E., Freedman, B. & Segev, I. Ion channel stochasticity may be critical in determining the reliability and precision of spike timing. Neural computation 10, 1679–1703 (1998).

20 Blazquez-Llorca, L. et al. High plasticity of axonal pathology in Alzheimer’s disease mouse models. Acta neuropathologica communications 5, 1–18 (2017).

21 Burke, R. E. & O’Malley, K. Axon degeneration in Parkinson’s disease. Experimental neurology 246, 72–83 (2013).

22 Suzuki, N., Akiyama, T., Warita, H. & Aoki, M. Omics approach to axonal dysfunction of motor neurons in amyotrophic lateral sclerosis (ALS). Frontiers in Neuroscience 14, 194 (2020).

23 Forsythe, I. D. Direct patch recording from identified presynaptic terminals mediating glutamatergic EPSCs in the rat CNS, in vitro. The Journal of physiology 479, 381–387 (1994).

24 Peterka, D. S., Takahashi, H. & Yuste, R. Imaging voltage in neurons. Neuron 69, 9–21 (2011).

25 Grienberger, C. & Konnerth, A. Imaging calcium in neurons. Neuron 73, 862–885 (2012).

26 Müller, J. et al. High-resolution CMOS MEA platform to study neurons at subcellular, cellular, and network levels. Lab on a Chip 15, 2767–2780 (2015).

27 Bullmann, T. et al. Large-scale mapping of axonal arbors using high-density microelectrode arrays. Frontiers in cellular neuroscience 13, 404 (2019).

28 Icha, J., Weber, M., Waters, J. C. & Norden, C. Phototoxicity in live fluorescence microscopy, and how to avoid it. BioEssays 39, 1700003 (2017).

29 Bakkum, D. J. et al. The axon initial segment is the dominant contributor to the neuron’s extracellular electrical potential landscape. Advanced biosystems 3, 1800308 (2019).

30 Radivojevic, M. et al. Electrical identification and selective microstimulation of neuronal compartments based on features of extracellular action potentials. Scientific reports 6, 31332 (2016).

31 Jäckel, D., Frey, U., Fiscella, M., Franke, F. & Hierlemann, A. Applicability of independent component analysis on high-density microelectrode array recordings. Journal of neurophysiology 108, 334–348 (2012).

32 Obien, M. E. J., Hierlemann, A. & Frey, U. Accurate signal-source localization in brain slices by means of high-density microelectrode arrays. Scientific reports 9, 1–19 (2019).

33 Ballini, M., et al. A 1024-channel CMOS microelectrode array with 26,400 electrodes for recording and stimulation of electrogenic cells in vitro. IEEE journal of solid-state circuits 49, 2705–2719 (2014).

34 Gagniuc, P. A. Markov chains: from theory to implementation and experimentation. (John Wiley & Sons, 2017).

35 Cormen, T. H., Leiserson, C. E., Rivest, R. L. & Stein, C. *Introduction to algorithms*. (MIT press, 2022).

36 Gutin, G., Yeo, A. & Zverovich, A. Traveling salesman should not be greedy: domination analysis of greedy-type heuristics for the TSP. Discrete Applied Mathematics 117, 81–86 (2002).

37 Franke, F., Quian Quiroga, R., Hierlemann, A. & Obermayer, K. Bayes optimal template matching for spike sorting–combining fisher discriminant analysis with optimal filtering. Journal of computational neuroscience 38, 439–459 (2015).

38 Buzsáki, G. Neural syntax: cell assemblies, synapsembles, and readers. Neuron 68, 362–385 (2010).

39 Kole, M. H. et al. Action potential generation requires a high sodium channel density in the axon initial segment. Nature neuroscience 11, 178–186 (2008).

40 Hu, W. et al. Distinct contributions of Nav1. 6 and Nav1. 2 in action potential initiation and backpropagation. Nature neuroscience 12, 996–1002 (2009).

41 Lorincz, A. & Nusser, Z. Molecular identity of dendritic voltage-gated sodium channels. Science 328, 906–909 (2010).

42 Duflocq, A., Chareyre, F., Giovannini, M., Couraud, F. & Davenne, M. Characterization of the axon initial segment (AIS) of motor neurons and identification of a para-AIS and a juxtapara-AIS, organized by protein 4.1 B. BMC biology 9, 1–19 (2011).

43 Stein, R. & Pearson, K. Predicted amplitude and form of action potentials recorded from unmyelinated nerve fibres. Journal of theoretical biology 32, 539–558 (1971).

44 Waxman, S. G. Determinants of conduction velocity in myelinated nerve fibers. Muscle & Nerve: Official Journal of the American Association of Electrodiagnostic Medicine 3, 141–150 (1980).

45 Huxley, A. & Stämpfli, R. Evidence for saltatory conduction in peripheral myelinated nerve fibres. The Journal of physiology 108, 315 (1949).

46 Björklund, U., Persson, M., Rönnbäck, L. & Hansson, E. Primary cultures from cerebral cortex and hippocampus enriched in glutamatergic and GABAergic neurons. Neurochemical research 35, 1733–1742 (2010).

47 Lewandowska, M. K., Radivojević, M., Jäckel, D., Müller, J. & Hierlemann, A. R. Cortical axons, isolated in channels, display activity-dependent signal modulation as a result of targeted stimulation. Frontiers in Neuroscience 10, 83 (2016).

48 Myers, J. P., Santiago-Medina, M. & Gomez, T. M. Regulation of axonal outgrowth and pathfinding by integrin–ECM interactions. Developmental neurobiology 71, 901–923 (2011).

49 Uesaka, N., Hirai, S., Maruyama, T., Ruthazer, E. S. & Yamamoto, N. Activity dependence of cortical axon branch formation: a morphological and electrophysiological study using organotypic slice cultures. Journal of Neuroscience 25, 1–9 (2005).

50 Grubb, M. S. & Burrone, J. Activity-dependent relocation of the axon initial segment fine-tunes neuronal excitability. Nature 465, 1070–1074 (2010).

51 Kumar, S. S. et al. Tracking axon initial segment plasticity using high-density microelectrode arrays: A computational study. Frontiers in neuroinformatics, 105 (2022).

52 Kuba, H., Oichi, Y. & Ohmori, H. Presynaptic activity regulates Na+ channel distribution at the axon initial segment. Nature 465, 1075–1078 (2010).

53 Ronchi, S. et al. Electrophysiological phenotype characterization of human iPSC-derived neuronal cell lines by means of high-density microelectrode arrays. Advanced biology 5, 2000223 (2021).

54 Hales, C. M., Rolston, J. D. & Potter, S. M. How to culture, record and stimulate neuronal networks on micro-electrode arrays (MEAs). JoVE (Journal of Visualized Experiments), e2056 (2010).

55 Yger, P. et al. A spike sorting toolbox for up to thousands of electrodes validated with ground truth recordings in vitro and in vivo. Elife 7, e34518 (2018).

56 Ronchi, S. et al. Single-cell electrical stimulation using CMOS-based high-density microelectrode arrays. Frontiers in neuroscience 13, 208 (2019).

57 Wagenaar, D. A., Pine, J. & Potter, S. M. Effective parameters for stimulation of dissociated cultures using multi-electrode arrays. Journal of neuroscience methods 138, 27–37 (2004).

